# Pervasive and dynamic release of Cryptic Genetic Variation in *Chironomus riparius*: Rethinking adaptation in fluctuating environments

**DOI:** 10.1101/2025.07.01.661462

**Authors:** Markus Pfenninger, Maria-Esther Nieto-Blazquez, Tilman Schell, Maximilian Geiss, Burak Bulut, Linda Eberhardt, Barbara Feldmeyer

## Abstract

The interplay between phenotypic plasticity and cryptic genetic variation (CGV) is crucial for understanding adaptation, yet the prevailing paradigm suggests CGV is primarily exposed under novel or extreme conditions. By examining gene expression responses along a natural temperature gradient in *Chironomus riparius*, we challenged this view. We found that the vast majority of expressed genes (63%) exhibit dynamic CGV, where interindividual expression variability scales continuously with distance from the selectively optimal temperature, a pattern also observed in higher-level traits like mutation rate and ROS levels. Genes with lower overall expression levels were less temperature-regulated, and thermal reaction norm shapes varied with gene function.

Unexpectedly, thermally plastic genes were more pleiotropic, often acting as hub genes, while CGV in gene expression was associated with lower pleiotropy. This pattern, and the observed strong recurrent selection on plastic genes with CGV, aligns with *C. riparius*’s adaptation to its highly fluctuating environment through selective tracking. We propose that this continuous, dynamic release of genetic variation is a necessary and inherent outcome of the polygenic nature of traits. This model fundamentally reshapes our understanding of adaptation, implying that populations can gradually and continuously adapt without requiring harsh conditions to expose hidden diversity. This leads to smoother adaptive landscapes, enhancing rapid adaptation and facilitating evolutionary innovation in the face of ongoing environmental change.

## Introduction

All organisms must cope with environments that fluctuate endlessly across all temporal (from minutes to centuries) and spatial (from centimeters to thousands of kilometers) scales to maintain a positive fitness (Bell, 2010). Broadly, these responses fall into two main categories: phenotypic plasticity and evolutionary adaptation. Phenotypic plasticity allows a single genotype to produce different phenotypes in varying environments, buffering against expected environmental variance within an individual’s lifetime and maintaining positive fitness (Pigliucci, 2001). Conversely, adaptation involves transgenerational genetic changes that enable populations to track and cope with environmental shifts over multiple generations (Rudman et al., 2018).

The capacity for both plasticity and adaptation is ultimately rooted in the underlying genetic variation within a population. A key mechanism driving phenotypic diversity and evolutionary potential are genotype-environment (G × E) interactions, where different genotypes exhibit distinct phenotypic responses across a range of environments. While G × E interactions are widely documented, their presence and magnitude vary considerably across populations and traits, and their full implications for how organisms respond to environmental change are still being uncovered (Saltz et al., 2018). In this context, cryptic genetic variation (CGV) plays a crucial role. CGV refers to genetic variation that remains unexpressed in the phenotype under certain conditions but becomes apparent under others (Paaby & Rockman, 2014). Mentioned as early as 1941 (Dobzhansky, 1941), CGV represents a hidden reservoir of potential phenotypic variation. This variation can be critical for evolutionary responses, especially when populations encounter novel or challenging conditions (Zheng et al., 2019). The existence of CGV is attributed to selection acting on robustness or canalization under stable conditions, which can maintain underlying genetic variation that is then released when environmental contexts change (Flatt, 2005).

Understanding how organisms respond to temperature fluctuations is particularly critical given the fundamental impact of temperature on biological processes (Knapp & Huang, 2022). The non-biting midge *Chironomus riparius* serves as an excellent model to explore these responses. The species is multivoltine, with up to 15 generations per year (Oppold et al., 2016). Larvae of *C. riparius* inhabit the sediments of shallow running waters, where both short-term solar radiation changes and air temperature shifts can cause strong temperature fluctuations (Leach et al., 2023) within and among generations. These transient spatial and temporal temperature variations significantly impact their fitness (Foucault et al., 2018; Nemec et al., 2013).

We addressed the organism’s response to naturally encountered temperature variation by investigating genome-wide gene expression levels. Being the most basal trait level, gene expression is a highly complex trait that integrates environmental cues with genetic background and directly influences all downstream phenotypes, making it an ideal candidate for studying thermal responses (Albert et al., 2018; Võsa et al., 2021). Previous research provided compelling evidence for CGV in temperature-regulated gene expression. For instance, in *Caenorhabditis elegans*, regulatory expression quantitative trait loci (eQTL) are often highly environment-specific and cryptic, with a significant portion induced only under specific temperature conditions like heat stress, indicating that CGV is largely determined by environmentally responsive regulatory elements (Snoek et al., 2017).

Similarly, in *Drosophila melanogaster*, extreme temperatures lead to significant allelic differences in gene expression, suggesting that temperature stress can uncover cryptic genetic variation by affecting gene expression regulation (Chen et al., 2015a). While temperature changes can alter gene expression widely, some studies, such as in the purple sea urchin, show effective buffering within gene networks, implying that CGV may not always be overtly expressed (Runcie et al., 2012). These studies collectively highlight the expectation for CGV in temperature-regulated gene expression and that these variations are often environment-specific and revealed under stress conditions.

However, a critical research gap remains in our understanding of how CGV is released along a continuous environmental gradient. It is commonly assumed that CGV is primarily released under not regularly experienced, extreme, or novel environmental conditions (Paaby & Rockman, 2014). Most previous studies on CGV have therefore focused on comparing “normal” versus “extreme” conditions (Walter et al., 2023), offering a limited view of its manifestation across a natural, continuous gradient. This experimental design restricts our ability to discern if CGV is an abrupt, threshold-dependent phenomenon or if it increases gradually with deviation from optimal conditions.

To address this gap, our study investigates gene expression responses in *C. riparius* across a natural temperature gradient, challenging the prevailing “extreme conditions only” view of CGV release.

Specifically, we ask:

- Which genes exhibit gene expression plasticity, and what are their population reaction norm shapes in response to a natural temperature gradient?
- Can we detect CGV in gene expression along this natural gradient, and if so, what “reaction norm” does it follow?
- What is the relation between gene expression plasticity and CGV, and what role does the gene’s pleiotropy play in this interaction?
- Does the evolution of gene expression plasticity and CGV align with the thermal environment’s fluctuations that natural populations of *C. riparius* experiences?

By answering these questions, our study aims to provide novel insights into the interplay between phenotypic plasticity, cryptic genetic variation, and pleiotropy at the fundamental level of gene expression. This mechanistic understanding is crucial for predicting organismal responses to environmental change, particularly in the context of climate change (Ledón-Rettig et al., 2014), and for elucidating how populations maintain evolutionary potential (McGuigan & Sgro, 2009) in dynamic environments.

## Material and Methods

### Experimental set-up

L3-stage *Chironomus riparius* larvae were taken from a laboratory culture, originally established with individuals from a population in a small stream (Hasselbach, Hessen, Germany) and regularly refreshed from this wild stock (Foucault et al., 2019). For each exposure temperature (5, 10, 15, 20, and 25 °C), 45 L3 larvae were maintained in aerated glass bowls with sand and medium, feeding *ad libitum*, as described (Foucault et al., 2019). These temperatures approximate the source habitat’s annual thermal range. The habitat’s mean water temperature during the primary reproductive period (May-September, encompassing most generations) is ∼16 °C (Waldvogel & Pfenninger, 2021), which approximately also matches the laboratory’s culture temperature. Larvae underwent temperature exposures in climate cabinets. To minimize developmental variation among treatments, exposure durations were inversely related to temperature: 2 days at 25 °C, 3 days at 20 °C, 4 days at both 15 °C and 10 °C, and 5 days at 5 °C. Per treatment, a total of 11-12 larvae were collected from the bowls, directly snap-frozen on dry ice, and stored at −80°C until further processing. Per treatment, a total of 11-12 larvae were collected from the bowls, directly snap-frozen on dry ice, and stored at −80°C until further processing.

### Gene Expression Analysis

Larvae were dissected on a −80°C gel pad. Three middle body segment were cut out for DNA extraction, the remainder of the body was submerged in 200µl TRIzol (Thermofisher). RNA was extracted using the Direct-zol RNA Miniprep Kits (Zymo Research) according to the manufacturer’s manual. Library preparation and sequencing of each of the 58 individuals was conducted at Novogene on a NovaSeq X Plus instrument.

Reads were aligned to the *C. riparius* reference genome v4 (Pettrich et al., 2024) using HISAT2 (v. 2.1.0) (Kim et al., 2019). Subsequently, the SAM files were sorted and converted to BAM format using Samtools (v. 1.20) (Li et al., 2009). The read counts were extracted using FeatureCounts (version 2.0.6) (Liao et al., 2014). We subsequently normalized the raw counts using DESeq2 (v. 3.19 (Love et al., 2014) and excluded genes with less than 10 read counts in at least eight individuals.

### Thermal reaction norms of gene expression

To characterize population thermal reaction norms of gene expression, we first determined whether genes were differentially expressed in at least one temperature comparison by used using (`scipy.stats`; Virtanen et al., 2020) between the two temperatures yielding each gene’s minimal and maximal mean normalized read counts. For such thermally plastic genes, expression data were fitted to six mathematical functions (linear, quadratic, logistic, exponential, beta, Michaelis-Menten) using Python `scipy.optimize` module (Virtanen et al., 2020). The best-fit model for each gene was selected based on the lowest Akaike Information Criterion (AIC) value. Estimated parameters from the best-fit model determined the direction of the response (e.g., linear increase or decrease with increasing temperature). Genes showing no significant difference were classified as “robust” (no detectable temperature response). These analyses were implemented in a custom Python script (Supplemental Script 1).

We tested with a permutation Runs-Test, whether temperature regulated genes with the same thermal reaction norms appeared spatially clustered in the genome. An observed number of runs smaller than expected by chance indicates spatial clustering, while a larger value is indicative of over-dispersion. We compared the observed number of runs with a random null-distribution obtained from 100,000 permutations. We compared the mean length of the runs for the different observed reaction norms with an ANOVA and Tukey’s post-hoc test. Both analyses were implemented in a custom Python script (Supplemental Script 2).

### Detecting cryptic genetic variation in gene-expression

We investigated two hypotheses regarding the relationship between environmental temperature and cryptic gene variation (CGV). First, we hypothesized that CGV becomes visible under the extreme experimental temperature conditions (5°C and 25°C) compared to intermediate temperatures (10°C, 15°C, 20°C). This was tested by performing a one-sided t-test comparing the mean standard deviation of gene expression at extreme temperatures against that at intermediate temperatures. Second, we hypothesized that gene expression variation scales positively with the distance to the selective optimum temperature. To test this, for each gene, we fitted the inter-individual standard deviations of gene expression at each temperature to the distance from the selective optimum temperature using a linear regression. We defined 16°C as the selective optimum, as this was the water temperature at which midges were sampled in the field and represents the mean summer water temperature at the sampling site (Waldvogel & Pfenninger, 2021). We used Akaike Information Criterion (AIC) to compare the fit of the observed data to each hypothesis. Genes exhibiting higher expression variation at intermediate temperatures or a negative slope in the linear regression were classified as not showing CGV under these environmental conditions and thus considered canalized. The same analytical approach was applied to the standard deviations of individual scores on gene-expression PCA axes that explained more variance than expected from a broken-stick model.

### Higher Trait Analysis

To infer whether trait variance scales with distance to the selective optimum along a temperature gradient in higher level traits, we analysed previously published data on ROS level (Bulut et al., 2025) and mutation rate (Waldvogel & Pfenninger 2021) in *C. riparius.* A measure of variance was fitted against the deviation from the selective optimum in a Bayesian framework (Bååth, 2014).

### Gene ontology analysis

The Interproscan database (v. July 2024) was used to obtain Gene Ontology (GO) information for all annotated genes in the *C. riparius* genome. A functional enrichment analysis in the category “biological function” was performed with various sets (see below). The analyses were conducted with the package topGO v.2.24.0 (Alexa & Rahnenführer, 2016) using the ‘parentchild’ algorithm and a p-value cut-off of <0.05 (Fisher’s test).

### Inference of network connectivity as a proxy for pleiotropy

The connectivity of a gene within a co-expression network is a useful proxy for its pleiotropy, as highly connected genes are more likely to influence multiple phenotypes. This relationship is observed across various studies, highlighting the role of gene network connectivity in understanding pleiotropic effects (Hämälä et al., 2020; Yuan et al., 2020). We used Weighted Gene Co-expression Network Analysis (WGCNA(Langfelder & Horvath, 2008)) to construct the network and the *intramodularConnectivity* function to infer kTotal as connectivity of each gene based on its r-values to all other genes in the whole network. To infer the relation between pleiotropy, temperature regulation and CGV and estimate their effects, we used a Bayesian implementation of a Two-way ANOVA (Bürkner, 2017) with kTotal as fixed effect and treatment temperature and expression variation as explanatory variables.

### Evolutionary parameters of expressed genes

To infer the evolutionary forces acting on the expressed genes, we referred to published data (Nieto Blazquez et al., 2025 for theta) or inferred them for this study (Tajima’s D) using *Variance-sliding.pl* from PoPoolation v.1.2.2 (Kofler et al., 2011). The estimates for the evolutionary parameters were calculated in 10 kb windows for a poolSeq data set gained from the natural population the experimental individuals were initially sampled from (NP0 sample from Pfenninger & Foucault (2022), Hasselbach, Hessen, Germany). If a gene fell completely into a single window, the respective evolutionary parameter value was taken, if it extended over more than a single window, their parameters were averaged. We analysed the relation between the parameters, temperature regulation and CGV using a Bayesian Two-way ANOVA as described above.

### CGV and plasticity of genes under fluctuating selection in a natural population

Based on the results from Pfenninger & Foucault (2022), which detailed selective tracking in response to environmental changes in the Hasselbach population, we extracted single nucleotide polymorphisms, whose temporal allele frequency trajectories were strongly correlated to water temperature changes (r > 0.7 or r < −0.7). For each of these SNPs, we inferred the closest gene. If this gene was expressed in the current data set, we extracted information on CGV and plasticity. We tested whether the proportion of genes showing both CGV and plasticity in this set differed from all other genes with a Bayesian test of proportions (Bååth, 2014).

### Population Genomic Simulation

We developed a simple forward-in-time, individual-based population genetic simulation using Python (version 3.9, Supplementary Script 3) to investigate the dynamics of quantitative genetic variation and phenotypic responses under varying environmental conditions and selection pressures under the condition that each allele has its own phenotypic reaction norm curve. This simulation builds upon the model described in Pfenninger (2025).

For each quantitative trait locus (QTL), allelic response curves were generated to define the contribution of each allele (A and a) to the phenotypic trait across a range of environmental conditions. These curves were linear, with randomly drawn intercepts and slopes for allele A, ensuring that allelic contributions remained within realistic bounds (0 to 1) across the defined environmental range. The allelic response curve for allele a was generated by randomly and subtly modifying the values drawn for allele a.

The initial population consisted of 1,000 adults. Genotypes at each QTL were assigned based on allele frequencies randomly drawn from a beta distribution (Beta (0.5, 0.5)), constrained between 0.1 and 0.9. The initial selective optimum for the trait was set to the mean phenotypic value of this nascent population, perturbed by a small Gaussian noise.

The simulation proceeded for 30 generations through discrete, non-overlapping generations. Reproduction involved random mating of adult individuals, producing a larger pool of juvenile offspring (10 times the adult population size). The trait was coded by a varying number of genes (20, 120, or 220). Offspring genotypes were determined by Mendelian inheritance, and their phenotypic values were calculated based on their genotype and the predefined allelic response curves at the current selective optimum. Hard selection was applied to the juvenile population, with survival probability exponentially decreasing with the deviation of an individual’s phenotype from the selective optimum. The strength of selection was expressed as an exponent in the exponential decline function, with values of 0.2, 0.5, or 0.8.

After the specified number of generations, a sample of 30 individuals was drawn from the final adult population. For each sampled individual, phenotypic values were calculated across the entire range of simulated environmental conditions. Population-level CGV was assessed by calculating the mean and standard deviation of phenotypic values across all sampled individuals for each environmental condition.

A linear regression was performed on the relationship between the standard deviation of the phenotypic trait and the absolute deviation from the selective optimum across environmental conditions. The slope, r value, and P-value from this regression, along with the population’s relative mean phenotypic distance to the selective optimum, were recorded for each simulation replicate (100 replicates per parameter combination).

## Results and Discussion

By examining changes in gene expression along a natural temperature gradient, we explored the intricate interplay between phenotypic plasticity and cryptic genetic variation (CGV), as well as their evolutionary implications. This approach enabled us to challenge the prevailing paradigm that CGV is primarily exposed under novel or extreme conditions.

### Genes with lower expression levels are less temperature regulated

We found that genes with lower expression levels are less regulated by temperature. Of the approximately 17,200 annotated genes in the *C. riparius* genome, 12,208 (71%) were expressed in the 58 larvae and met the criteria for analysis. Among these expressed genes, 7,811 (64%) exhibited a plastic response, demonstrated by significant expression differences between at least two temperatures. In contrast, the expression of 4,397 (36%) genes remained unaffected by temperature variations within the typical physiological range, indicating robustness. The overall mean expression level of a gene significantly influenced its classification as plastic or robust (Supplemental Figure 1, log difference = 0.42, 95% HDI 0.39-0.45, pp 100%, effect size = 0.58). Specifically, genes with lower average expression were more frequently classified as robust, a pattern previously observed (Pallares et al., 2023; Wolf et al., 2023). One possible explanation is that genes with low expression levels may play more specialized or context-dependent roles or may be less critical for cellular function, thus not requiring dynamic regulation (Inoue & Horimoto, 2017; Romero et al., 2012). Consequently, their expression might be more constitutive and less responsive to environmental changes. However, it remains unclear whether the observed moderate effect stems from a biological mechanism or if the statistical detection of expression differences is less probable at lower expression levels.

### Thermal gene expression reaction norms are variable and associated to their function

Among the temperature-dependent genes, the thermal reaction norms of 1,954 genes were best described by a linear model (25% of all regulated genes), with 990 increasing and 964 decreasing in expression with temperature. Gene Ontology (GO) terms associated with a linear increase in gene expression with temperature were enriched for core functions in energy production, biosynthesis, and macromolecular turnover (Supplemental List 1A). This likely reflects a coordinated cellular response to maintain optimal function, cope with increased metabolic demands (Tattersall et al., 2012), and adapt to stress associated with higher temperatures (Bulut et al., 2025). Conversely, genes showing a negative linear response to temperature were enriched for functions in the negative regulation of metabolism and biosynthesis (Supplemental List 1B). This may represent a cellular strategy to slow down non-essential or energetically costly processes and reallocate resources when faced with increasing thermal stress (O’Connor & Bernhardt, 2018), or it may simply reflect the general increase of chemical reaction rates with temperature, requiring fewer enzymes to maintain the same activity level (Jaquet et al., 2022).

1787 genes showed a quadratic expression pattern (21.2%, 1051 upwards highest expression at extreme temperatures, 736 downwards, highest expression at intermediate temperatures). The genes higher expressed at both cold and hot extreme temperatures were enriched for functions in transport, regulation of programmed cell death, stress response and homeostasis (Supplemental List 1C), indicating a targeted plastic reaction to suboptimal thermal conditions (Somero, 2020). The inverse response, a higher gene expression at temperatures closer to the adaptive optimum encompassed functions like biosynthesis of core building blocks, protein localization and assembly, protein quality control, regulation of gene expression and translation. The increased expression of these genes indicated robust growth, efficient synthesis of cell building blocks, and the precise assembly of complex cellular structures under optimal conditions. Their downregulation at temperature extremes suggested that these processes are either highly sensitive to temperature fluctuations, or they are actively suppressed because they are too energy-intensive, too prone to error, or not essential for immediate survival under harsh conditions (Grigaitis & Széliová, 2023).

With 2275 reaction norms the largest set was adequately described by an exponential equation (30.8%, 1350 increase, 925 decrease, Figure 1A). GO-term enrichment indicated that the functions of the such regulated genes were in positive regulation of growth, biosynthesis, and cell cycle, lipid metabolism and carbohydrate and amino sugar metabolism/catabolism (Supplemental List 1E).

**Figure 1.**
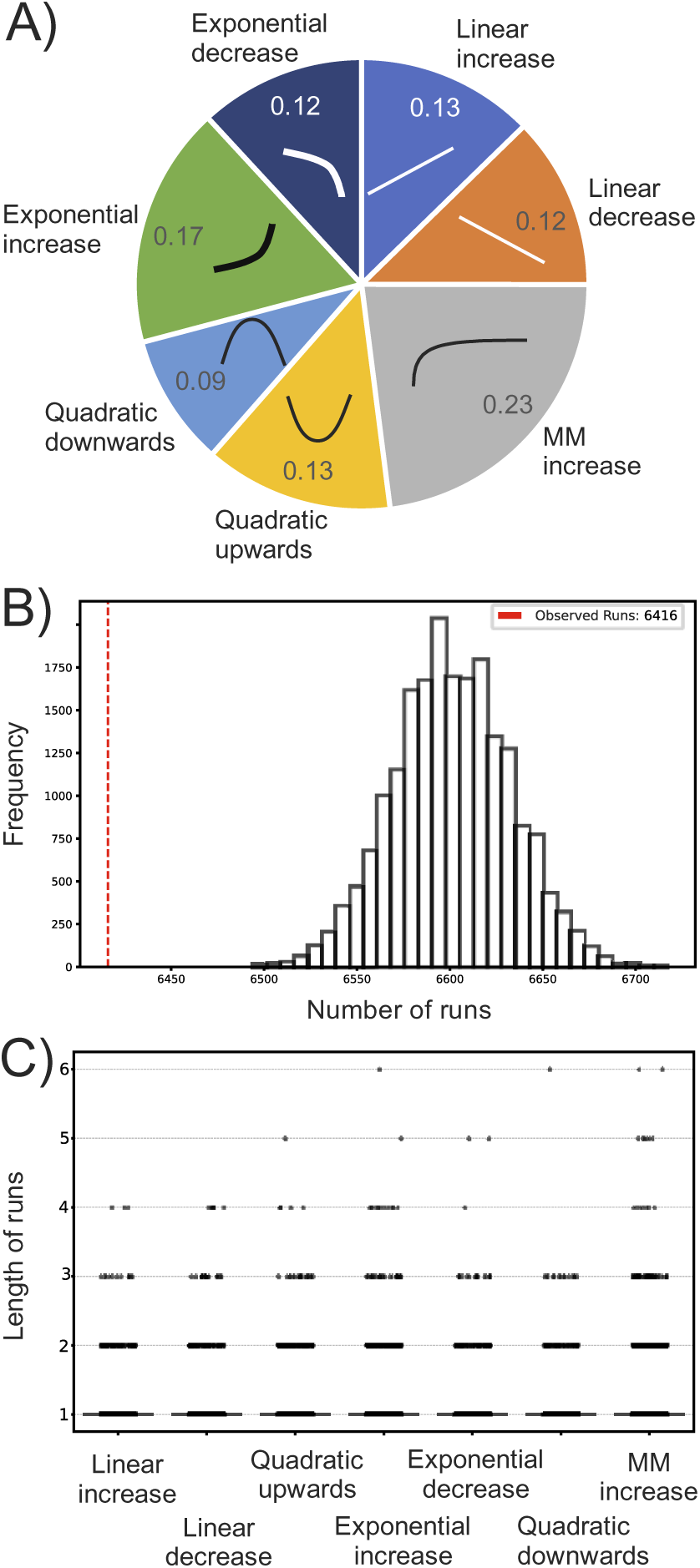
Thermal reaction norms of expressed genes and spatial co-regulation. A) Frequency of the observed reaction norms. B) Comparison of the observed runs of genes following the same reaction with simulated random expectations. C) Distribution of length of runs.

Depending on the threshold temperature for the strong increase, this either suggested maximal metabolic activity, rapid cell division, and efficient resource utilization, or a stressful but still survivable threshold, where the midges might initiate a vigorous and rapid adaptive response to cope with the heat. This included bolstering protein quality control, adjusting membrane composition, and potentially accelerating certain metabolic pathways to manage the stress effectively (Balogh et al., 2013; Lee et al., 2016). The corresponding exponential negative response to temperature suggested a rapid shutdown or suppression of gene expression. This might induce syndromes like reduced cellular transport and traffic, decreased biosynthesis of complex molecules, RNA metabolism downregulation, mitochondrial function decline and DNA repair compromise. These functions indicated that the threshold to a state of severe stress, damage control, or even cellular demise/quiescence was passed. The organism was rapidly and drastically shutting down complex, energy-intensive, and dynamic processes to conserve energy and prevent further damage or even reflect beginning cellular breakdown (Adjirackor et al., 2020).

With 1795 reaction norms, a substantial proportion corresponded to a Michaelis-Menten model (23.0%, only increase found), a response model not often described for gene expression (Liu et al., 2017). This enzyme-like kinetics suggested that the expression of these genes is limited by the “saturation” of a key component. This interpretation is strengthened by one of the major themes of the enriched GO-terms, regulation of catalytic activity. Other major themes, like endocytosis, DNA recombination and double-strand break repair, suggested that the transcription, albeit temperature dependent, might plateau due to the saturation of the processing cellular machinery, because maximum production capacity is reached or lacking availability of necessary components. Reaction norms resembling a beta function, often found for fitness traits, were not observed.

A similar study in *Drosophila melanogaster* found broadly comparable results regarding thermal gene expression patterns (Chen et al., 2015b). However, direct comparison is challenging due to key methodological differences: Chen et al. examined a smaller temperature gradient (13-29°C), reported a lower proportion of expressed genes (53% vs. 71% in our study), and restricted their reaction norm classification to only linear or quadratic models. Nevertheless, the general trends in enriched gene functions associated with thermal response, particularly for linear and quadratic patterns, were similar to those observed here (Chen et al., 2015b), suggesting potentially conserved cellular strategies to temperature variation across diverse insect taxa.

### Spatial Proximity Enhances Co-Regulation

We observed that spatially consecutive genes tended to share the same reaction norm shape significantly more frequently than expected by chance (Runs test, N runs observed = 6416, p < 0.001; Figure 1B). The different reaction norms did not contribute equally to this pattern (ANOVA, F = 14.77, p = 7.9 x 10^-17^); genes with Michaelis-Menten-like and, to some extent, exponential increase reaction norms tended to have longer runs (Figure 1C, Supplemental Table 1). This finding aligns with other studies demonstrating that co-regulated genes often cluster. For instance, (Pannier et al., 2017) found that co-regulated genes within 1–20 kb of each other showed significantly higher co-expression levels, independent of shared transcription factors. One possible reason for this phenomenon is that chromatin domains with similar epigenetic states or regulatory programs tend to co-localize in the nucleus, enabling shared transcriptional machinery (Khrameeva et al., 2012).

### Cryptic genetic variation is pervasive and scales mostly linearly with distance to selective optimum

The expression patterns of a substantial proportion of genes, 7,712 (63%), were consistent with the release of cryptic genetic variation (CGV) along the thermal gradient. Among these, 774 expression patterns were better described by the extreme value model, while the interindividual expression variation of 6,968 genes increased linearly with the distance to the selective optimum (Figure 2A). This indicates that the expression of the vast majority of genes exhibited signs of CGV when individuals were exposed to temperatures deviating from the selectively optimal conditions.

**Figure 2.**
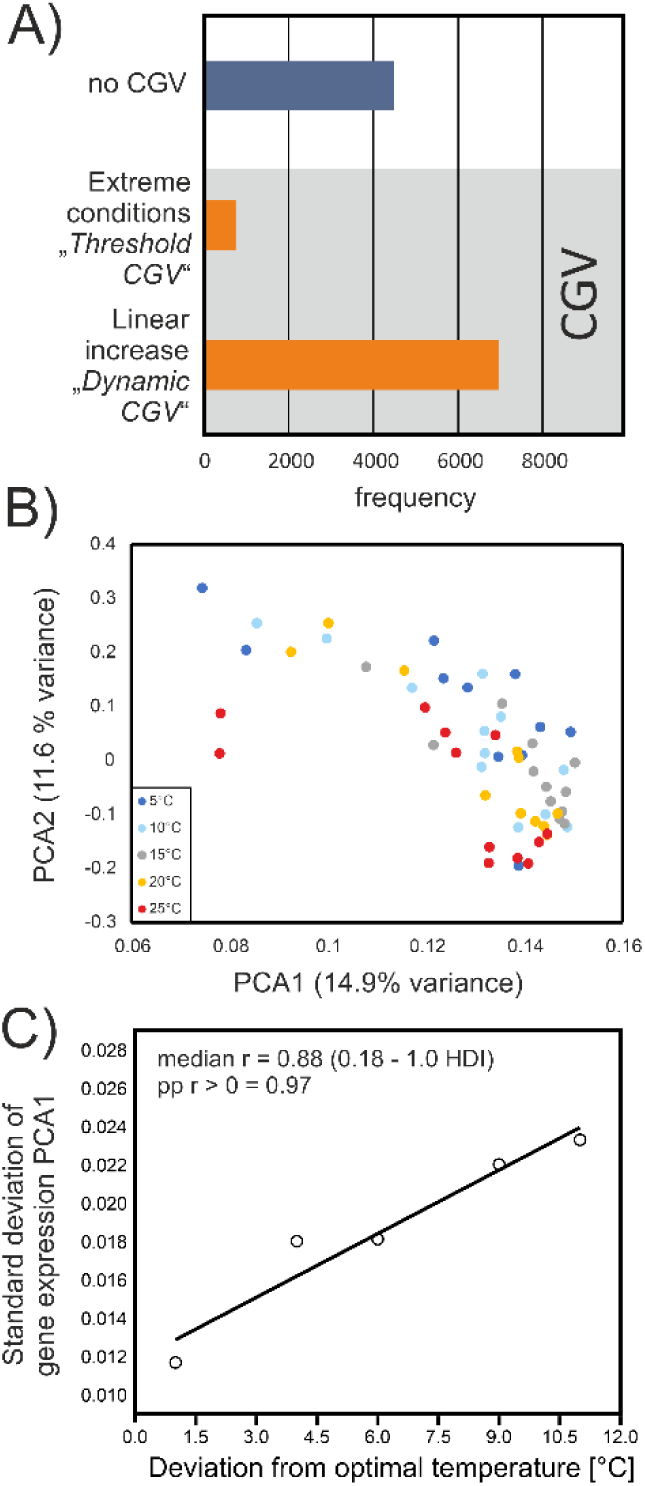
Cryptic genetic variation along a thermal gradient in the gene expression of *C. riparius*. A) Frequency of expression patterns for all expressed genes. B) Plot of the individual scores on PCA1 and PCA2 based on the gene expression of all genes. C) Correlation of the standard deviation of individual scores against the deviation from the optimal temperature.

Crucially, in most cases, this interindividual variability of gene expression increased linearly with distance from the optimal temperature, with relatively few instances showing CGV only under extreme conditions. To our knowledge, the observation and prevalence of such a linear, gradual relationship between CGV and distance to optimal conditions is novel. To distinguish this newly recognized mode of variable CGV release from the traditional threshold model, we henceforth term it Dynamic CGV. The lack of previous observations might be attributed to experimental designs in earlier CGV studies that focused primarily on “normal” versus extreme conditions and did not explore continuous environmental gradients.

The overall strength of gene expression had no major influence on whether a gene exhibited CGV. Although our analysis indicated a statistically significant difference, the effect size was marginal (difference of log10 means = 0.078, 95% HDI 0.048-0.11, posterior probability (pp) of difference > 0 = 100%, effect size = 0.099), suggesting no substantial biological relationship.

Principal component analyses (PCA) of gene expression data identified five axes explaining more variance than expected, together accounting for over 80% of the total variation (Figure 2B). Notably, the standard deviation of individual scores on PCA1 (representing 36.5% of the total variation) increased strongly with deviation from the optimal temperature (r = 0.93, p = 0.01, Figure 2C). This finding underscores that Dynamic CGV, as revealed by the broad increase in interindividual gene expression variability along the thermal gradient, is not a statistical artefact but represents a pervasive biological characteristic governing major aspects of gene expression in response to temperature. In contrast, the standard deviations of PCA axes 2-5 were uncorrelated with temperature deviations (Supplemental Table 2).

### Higher-Level Traits exhibit similar Dynamic CGV patterns

A linear increase of phenotypic variation with distance to the selective optimum temperature was also observed for higher-level phenotypic traits, including mutation rate (median r=0.60, 95% HDI 0.20-0.98, posterior probability of positive correlation (pp) = 0.90; Figure 3A, Waldvogel & Pfenninger, 2021) and internal reactive oxygen species (ROS) level (median r=0.71, 95% HDI = 0.07-0.98, pp=0.97; Figure 3B, Bulut et al., 2025). This strongly suggests that the Dynamic CGV pattern, predominant in gene expression, is a general mechanism that also characterizes and likely shapes cryptic genetic variation in more complex, higher-level traits. Similar observations concerning phenotypic variation in response to temperature gradients have been noted in other species and traits, such as age at pupation in *Aedes japonicus* (Reuss et al., 2018).

**Figure 3.**
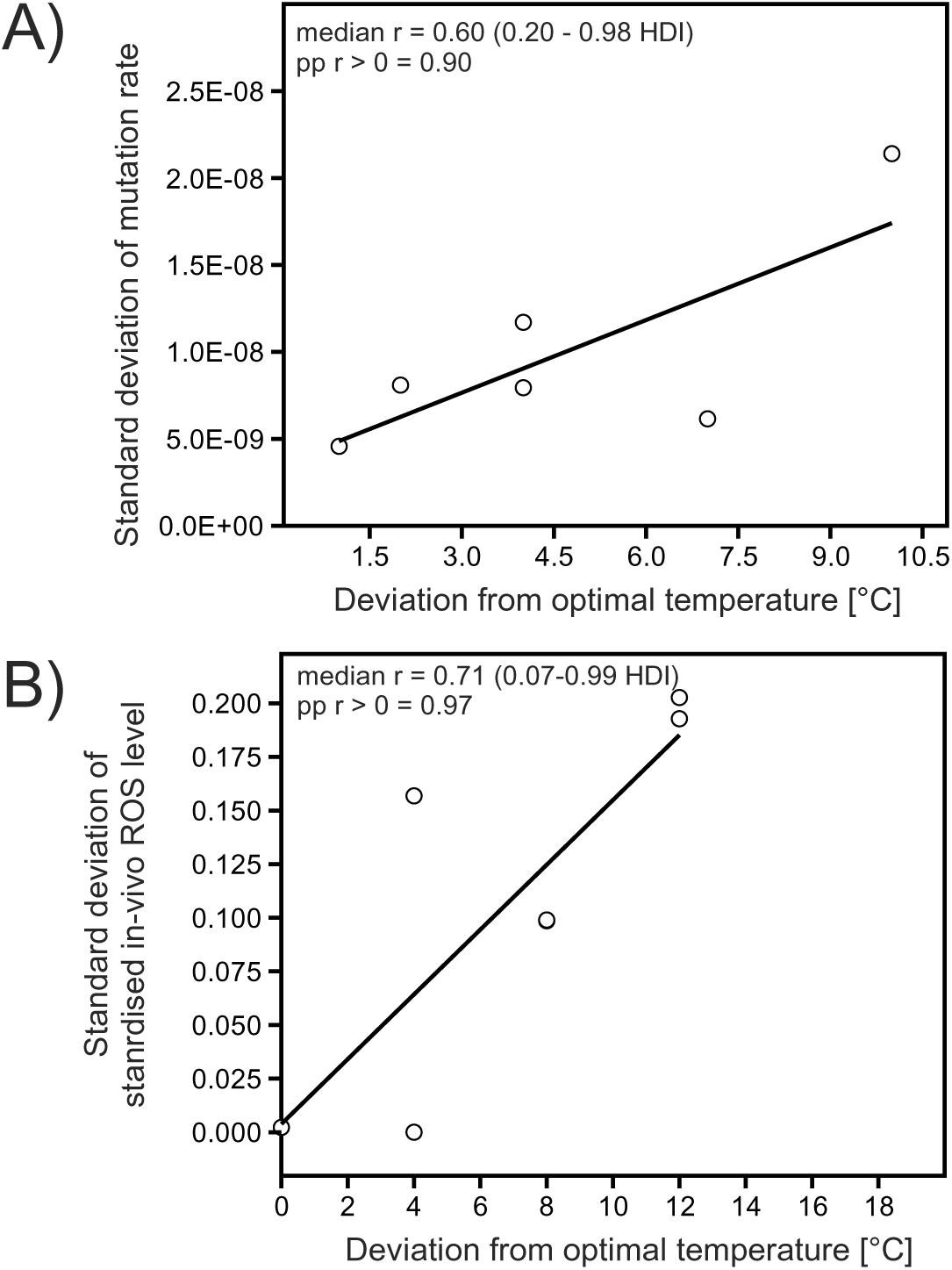
Relation between interindividual trait variance and deviation from fitness optimum temperature for higher level phenotypic traits. A) Standard deviation of mutation rate among individuals against deviation from optimal temperature (Waldvogel & Pfenninger, 2021), B) Standard deviation of *in-vivo* ROS level among individuals against deviation from optimal temperature (Bulut et al., 2025).

### The interplay of phenotypic plasticity and cryptic genetic variation shapes evolutionary potential and robustness

To understand the combined evolutionary implications of thermal plasticity and CGV, we categorized *C. riparius* genes into four distinct groups based on the presence or absence of thermal expression plasticity and CGV (Table 1).

**Table 1.** Cross table of gene expression classifications into plastic/robust and CGV/no CGV pattern.

| | plastic | robust | $\Sigma$ |
| --- | --- | --- | --- |
| CGV | 4516 | 2242 | 6758 |
| No CGV | 3295 | 2155 | 5450 |
| $\Sigma$ | 7811 | 4397 | $\Sigma\Sigma$ 12,208 |

The first category, comprising 4,516 thermally plastic genes exhibiting CGV, represents genes with significant evolutionary potential for thermal reaction norm evolution. Gene Ontology (GO) enrichment analyses revealed main themes of cellular maintenance, protein processing, and metabolic regulation, with a strong emphasis on cellular components and responses to stress (Supplemental List 1A). The temperature regulation and CGV in these processes are often interconnected, particularly for genes involved in cellular maintenance and stress responses (Camus et al., 2017), reflecting a critical need to maintain homeostasis under varying thermal conditions. The presence of CGV in these plastic genes provides a crucial mechanism for populations to adapt to both predictable (e.g., seasonal temperature shifts) and unpredictable environmental changes by revealing previously hidden genetic potential (Paaby & Rockman, 2014). Consequently, these genes are likely instrumental in the reported adaptive tracking of seasonal temperature changes in *C. riparius* (Pfenninger & Foucault, 2022). Their biological significance for the species was further supported by their statistical overrepresentation (median estimate of association difference = 6.8%, 95% credible interval = 4.9-8.2%, posterior probability of association = 100%).

The second set of genes, totalling 2,242 genes, were robust to temperature variation but nevertheless exhibited CGV in response to temperature. GO term analysis revealed enrichment for genes with a strong emphasis on anabolism and positive regulation of biosynthesis, particularly involving nitrogen-containing compounds and macromolecules (e.g., RNA and peptides). While fundamental to many biosynthetic pathways, the expression of these genes may have evolved for robustness to temperature fluctuations, ensuring a continuous supply of vital molecules across the typical physiological range. Alternatively, regulation might occur predominantly downstream (e.g., enzyme feedback inhibition, allosteric control), making upstream gene expression plasticity less critical (De Hijas-Liste et al., 2015).

The presence of CGV in these robust genes can be attributed to several mutually non-exclusive factors. Firstly, complex biosynthetic pathways often feature redundancy, alternative routes, or robust feedback loops (Bhalla & Iyengar, 1999). This inherent buffering capacity can mask the phenotypic effects of individual gene variants under suboptimal conditions, meaning natural selection may not consistently purge every slightly suboptimal variant if the overall pathway output maintains fitness. Secondly, in environments with strong temporal or spatial fluctuations, CGV in biosynthetic pathways could be advantageous; a previously “hidden,” neutral variant might become beneficial under specific conditions (Grieshop et al., 2024). Thirdly, pleiotropy can play a role: a gene involved in biosynthesis might have other, less critical functions, where transcriptional variation might be neutral or even beneficial under non-optimal conditions, despite being neutral or slightly deleterious for the primary pathway (Mackay & Anholt, 2024). Ultimately, even for robust genes, CGV provides untapped potential for expression evolution if thermal conditions change over extended evolutionary timescales.

The third group comprised 3,295 thermally plastic genes with canalised expression. The enriched GO terms for this group centred on themes of cellular internal management, resource allocation (e.g., amino acids), and the cell’s ability to self-regulate and respond to internal and external challenges to maintain homeostasis and viability (Supplemental List 1C). This strongly suggests that these genes play critical, unbuffered roles in response to temperature variation. Given their absolute essentiality for immediate cell viability and function across the physiological temperature range, even subtle gene expression variations in these pathways could prove immediately deleterious under both normal and fluctuating temperatures (Flatt, 2005). Thus, the optimal expression levels for these genes are likely precisely defined across various temperatures, with deviations generally incurring significant fitness costs.

The final category consisted of 2,155 genes that were robust to temperature changes and also canalised (Table 1). GO term analysis revealed their primary functions to be in developmental processes and the regulation of cell growth and differentiation, particularly within the nervous system. The expression of genes critical for these fundamental developmental processes must be robust against environmental fluctuations (Braendle & Félix, 2008). This robustness is typically achieved through mechanisms like redundant pathways, feedback loops, and molecular chaperones (e.g., HSP90), which collectively ensure a consistent developing phenotype. Indeed, the expression of developmental genes is often strongly canalized (Hornstein & Shomron, 2006), as any significant deviation or perturbation is likely to have severe fitness consequences and is thus under strong purifying selection.

### Thermally plastic genes are more pleiotropic, genes with cryptic genetic variation are less pleiotropic

As expected from the literature (Barbitoff et al., 2024; Singh & Yi, 2021), more pleiotropic genes— defined as genes with a higher number of gene expression network connections—tended to be, on average, more highly expressed (r=0.43, 95% HDI 0.41-0.45, posterior probability of positive correlation = 100%, Supplemental Figure 2). This association is consistent with the understanding that pleiotropic genes often engage in more protein-protein interactions and form more complex functional networks, necessitating higher expression levels (Singh & Yi, 2021).

Our Bayesian analysis revealed that thermally plastic genes are, on average, much more pleiotropic than genes whose expression is not regulated by temperature (Figure 4, median effect: +25.16 connections, 95% HDI 22.64 - 27.77, posterior probability of effect > 0 = 100%). Inspection of the raw data (Figure 4) indicated that this elevated mean pleiotropy is not uniformly distributed across all plastic genes, but rather driven by a restricted number of genes exhibiting a very high number of connections, likely acting as hub genes that control the expression of many other genes (Arnatkeviciute et al., 2021).

**Figure 4.**
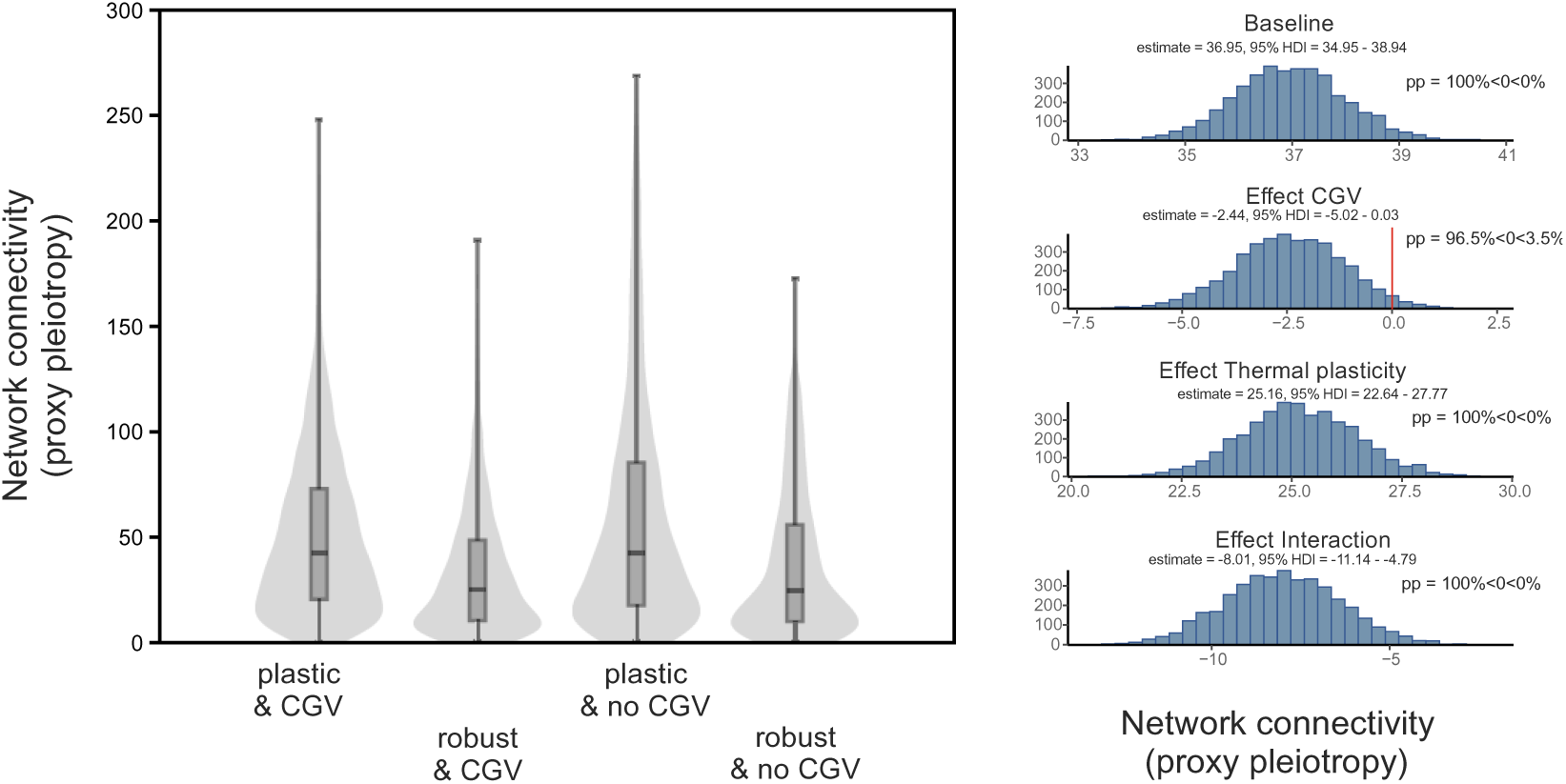
Effect of CGV and temperature regulation on a gene’s pleiotropy as estimated from it’s gene expression network connection. Data distribution in each category and Bayesian estimates of parameter effect size.

In contrast, the effect of cryptic genetic variation (CGV) on pleiotropy was slightly negative (median effect: −2.44, 95% HDI −5.02 - 0.03, posterior probability of effect < 0 = 97%). Even more notably, the interaction between temperature regulation (plasticity) and CGV resulted in a stronger negative effect on pleiotropy (median effect: −8.01, 95% HDI −11.14 - −4.79, posterior probability of effect < 0 = 100%, Figure 4).

These findings, particularly the higher pleiotropy of plastic genes and the lower pleiotropy associated with CGV, conflict with published hypotheses. Traditional views suggest that thermally regulated genes should exhibit less pleiotropy, as lower pleiotropy is thought to confer greater adaptability to temperature changes, making highly pleiotropic genes less flexible in their expression responses to thermal stress (Papakostas et al., 2014). Conversely, genes associated with CGV are often hypothesized to show increased pleiotropy, given their involvement in multiple traits and complex genetic interactions (Watanabe et al., 2019; Yadav et al., 2016).

However, this apparent discrepancy can be explained by the distinct regimes of environmental variance in which these organisms have evolved. The larvae of *C. riparius* inhabit the sediments of shallow running waters, where both short-term changes in solar radiation and air temperature can induce strong and rapid temperature fluctuations (Leach et al., 2023). Consequently, these midges frequently experience transient spatial and temporal temperature variations within a single generation (e.g., (Pfenninger et al., 2022)), which are known to significantly impact their fitness (Nemec et al., 2013). In such highly variable environments, coordinated plastic responses across numerous traits—often controlled by highly pleiotropic hub genes—could be highly beneficial for maintaining fitness. Conversely, CGV at loci with higher pleiotropy might prove more deleterious than beneficial for a larger proportion of individuals during frequent, transient temperature perturbations, leading to selection against such highly pleiotropic CGV in *C. riparius*.

### Selective regime on genes with temperature regulation and CGV is consistent with selective tracking

Inspecting classical evolutionary parameters revealed the selective regimes acting on the genes in the four categories. Compared to the entire genome, all expressed genes were located in genomic regions exhibiting lower theta in a natural population, indicating purifying selection (Figure 5A). While cryptic genetic variation (CGV) showed a minor tendency to slightly increase theta, and temperature regulation itself had a subtle negative effect, the interaction between temperature regulation and CGV exerted a substantial negative impact on theta. This strong reduction suggests that temperature-regulated genes with CGV are under intense, recurrent selection relative to other expressed genes (Charlesworth, 2009) (Figure 5A).

**Figure 5.**
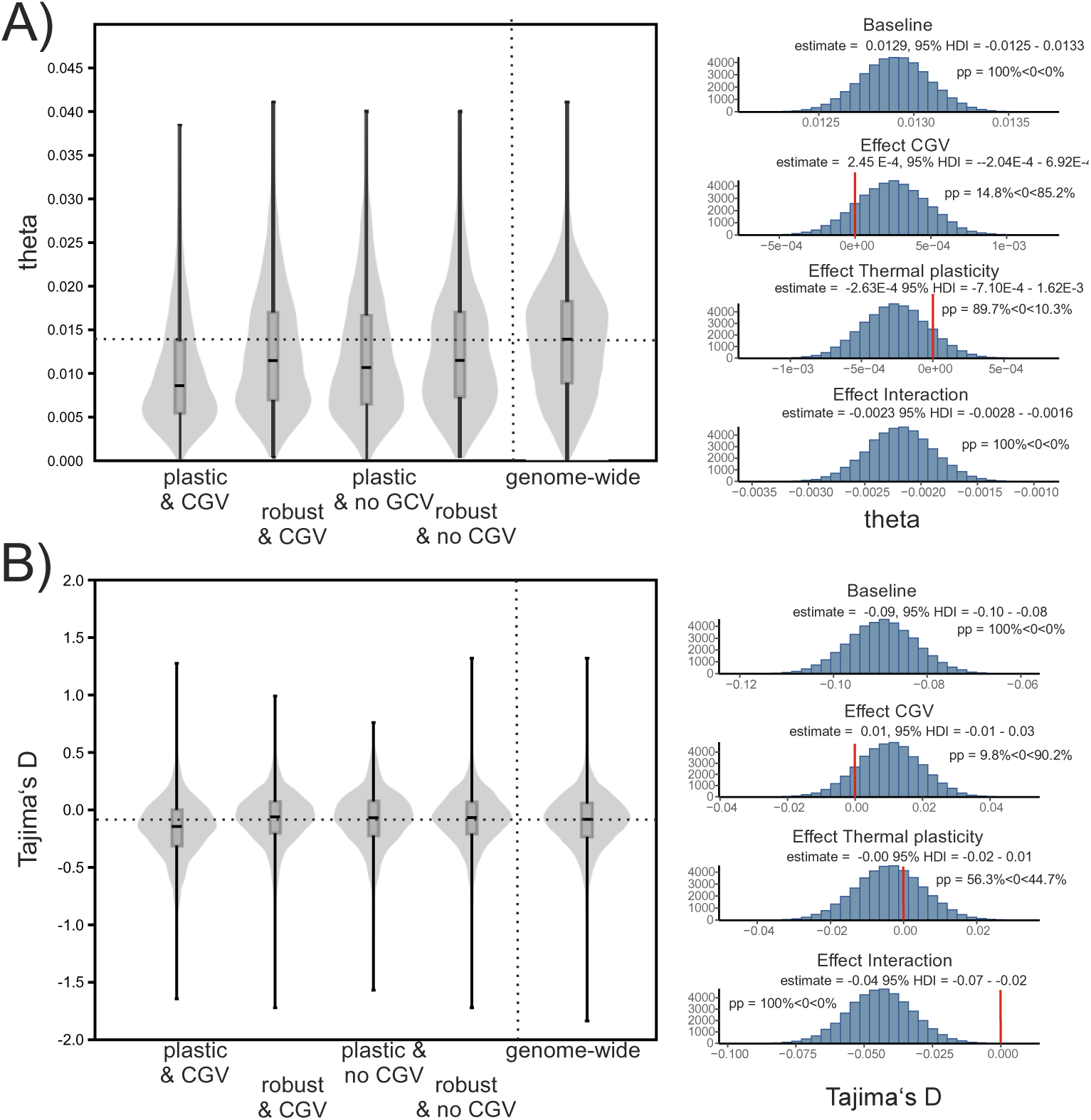
Effect of gene expression plasticity and CGV on evolutionary patterns. A) Effect on theta, B) effects on Tajima’s D. Both parameters were estimated for the regional genomic locations (10 kb windows) of the respective genes.

The site-frequency spectrum, measured as Tajima’s D, presented a complementary picture. While the mean of regions containing expressed genes did not deviate from the genome-wide average (Figure 5B), CGV showed a slightly positive effect on Tajima’s D. This positive value indicates a relative lack of rare alleles in these regions, which typically suggests balancing selection maintaining older alternative variants. In this context, it implies that CGV is associated with alleles contributing to different gene expression reaction norms across the temperature gradient. However, the interaction between plasticity and CGV, despite plasticity’s individually inconspicuous effect, tended to significantly lower Tajima’s D (Figure 5B). This reduction signifies stronger than average selection acting specifically on thermally plastic genes that also exhibit CGV (Charlesworth, 2009).

This pattern of relatively stronger selection on thermally plastic genes with CGV aligns well with the known selective regime of *C. riparius*. A large proportion of genes in this species are known to be under fluctuating selection, characterized by frequent shifts in the direction of selective pressures in response to seasonal or other rapid environmental fluctuations (Pfenninger et al., 2022; Pfenninger & Foucault, 2022). In natural populations, such fluctuating selection, as experienced by *C. riparius*, can repeatedly expose and favour different cryptic variants. This process not only maintains genetic diversity (Tuyishimire et al., 2025) but also enables rapid evolutionary responses, a mechanism often referred to as selective tracking (Ledón-Rettig et al., 2014; Nosil et al., 2024). Indeed, in a natural population, the genes close to fluctuating selected SNPs in response to temperature variation were enriched for both plastic and CGV genes by 7.8% compared to the entire gene set (95% HDI interval = 0.01 – 0.15, posterior probability of enrichment 1.4% <0 < 98.6%). This underscores the importance of CGV for the adaptive potential in this and probably other species. For example, studies of stick insects have demonstrated how negative frequency-dependent selection and environmental variability can drive predictable fluctuations in cryptic morph frequencies (Nosil et al., 2024).

Our results reveal pervasive CGV in the expression of genes that are not only robustly expressed across a wide range of temperatures but also appear to be under strong purifying and fluctuating directional selection. This finding significantly challenges traditional theoretical expectations, which generally posit that CGV should be more prevalent in genes underlying conditionally and rarely expressed traits. Under this relaxed selection scenario, mutations can accumulate in such genes or their regulatory regions without being exposed to purifying selection (Kawecki, 1994; Snell-Rood et al., 2010; Van Dyken & Wade, 2010). In contrast, the pervasive and dynamic release of CGV observed here, particularly for genes under fluctuating selection, aligns more closely with models suggesting that recurrently exposed CGV will have a reduced genetic load (Eshel & Matessi, 1998).

### Dynamic CGV as an inherent characteristic of polygenic traits

We therefore propose an alternative hypothesis for the observed release dynamics of cryptic genetic variation (CGV). At the fitness optimum, phenotypes of a population exhibit minimal variation in fitness-relevant traits due to strong purifying selection acting against deviations from the selective optimum. As noted by (Mather, 1943), these optimal phenotypes are necessarily similar under the specific environmental conditions for which they were selected. However, this phenotypic similarity often masks extensive underlying genetic diversity, as these selectively equivalent, polygenic phenotypes can be produced by numerous different genotypes through differential allelic contributions from many loci.

To illustrate this ‘many-to-one mapping’ principle, consider an intermediate phenotype of a quantitative trait determined by 10 equally contributing biallelic loci; this single phenotype can be encoded by 8,953 distinct genotypes. Crucially, each of these alleles, influenced by idiosyncratic gene-by-environment (GxE) and gene-by-gene (GxG) interactions (i.e., epistatic background), may possess its own allele-specific phenotypic response curve concerning its allelic contribution along an environmental gradient. When these genotypically diverse individuals are exposed to conditions deviating from the optimum, these inherent differential response curves manifest, leading to the production of increasingly variable phenotypes (Bürger & Lynch, 1995; Paaby & Rockman, 2014).

Our population genetic forward-in-time simulations demonstrate that the proposed mechanism consistently leads to the emergence of dynamic CGV. After 30 generations of simulated evolution, populations exhibited near-perfect adaptation to the selective optimum (mean relative deviation <0.001, 95% HDI =0.000–0.002). Importantly, approximately 99% of all 900 simulations revealed a significant positive correlation between the standard deviation of the trait and its deviation from the selective optimum, robustly confirming the dynamic CGV pattern. The strength of this dynamic response, quantified by the slope of the linear regression, was negatively influenced by stronger selection. Genetic architecture and its interaction with selection strength showed only minor (negative and positive, respectively) effects (Supplementary Figure 3). While these relatively simple simulations successfully validate that the hypothesized mechanism of alleles with idiosyncratic reaction norms can indeed produce dynamic CGV, it is crucial to note that they do not constitute definitive proof of its ultimate cause in natural systems.

We thus propose that dynamic CGV, as observed here, is a necessary and inherent outcome of the polygenic nature of many, if not most, quantitative traits. This perspective shifts the burden of explanation: under this hypothesis, phenomena such as threshold CGV or a complete absence of CGV would, in fact, require the evolution of specific buffering mechanisms to suppress the natural tendency for variation to be revealed when environmental conditions deviate from the selective optimum, a mechanism extensively explored by the traditional view (Paaby & Rockman, 2014).

### Conclusion

Our study reveals that phenotypic plasticity and cryptic genetic variation (CGV) pervasively shape gene expression responses to natural temperature gradients in *C. riparius*. Contrary to the prevailing paradigm that CGV is primarily exposed only under extreme or novel conditions, we demonstrate that the vast majority of expressed genes exhibit dynamic CGV, scaling continuously with their deviation from the selectively optimal temperature. This principle extends beyond gene expression to higher-level phenotypic traits, underscoring its broad generality. Furthermore, we uncovered an unexpected and crucial relationship: thermally plastic genes are more pleiotropic, whereas CGV in gene expression tends to be less pleiotropic—a pattern likely reflecting *C. riparius*’s adaptation to its highly fluctuating and transient habitat.

The continuous, dynamic release of genetic variation in changing environments, as elucidated by our findings, fundamentally reshapes our understanding of adaptation. It suggests that populations can achieve gradual and continuous adaptation without requiring harsh conditions to expose hidden diversity, thereby leading to smoother adaptive landscapes that facilitate rapid adaptation and selective tracking. We propose that this dynamic CGV model, by perpetually revealing novel phenotypic variation, represents a crucial yet under-appreciated mechanism for evolutionary innovation in the face of ongoing environmental change.

## Data Availability Statement

The genomic data underlying this article are available at the European Nucleotide Archive (ENA), and can be accessed under the accession number PRJEB89193. Normalised gene expression read counts, gene-wise parameters and GO-result tables can be found at Zenodo doi: 10.5281/zenodo.15729658.

## AI Disclaimer

Please note that in the generation of the manuscript, AI was extensively used to search scientific literature (consensus.ai, ask.ork.org, googleScholar), store and query it (NotebookLM). The Python-scripts created were debugged, corrected and annotated (codepal.ai, Gemini) and the final text was proofread for grammar and style (Gemini, Le Chat) by AI.

## Acknowledgments

No funding to report.

