## Supplementary Info for "Pervasive and dynamic release of Cryptic Genetic Variation in *Chironomus riparius*: Rethinking adaptation in fluctuating environments"

### Supplemental material

#### Part A Supplemental Figures

**Supplemental Figure 1**. Log expression level of temperature plastic versus robust genes.

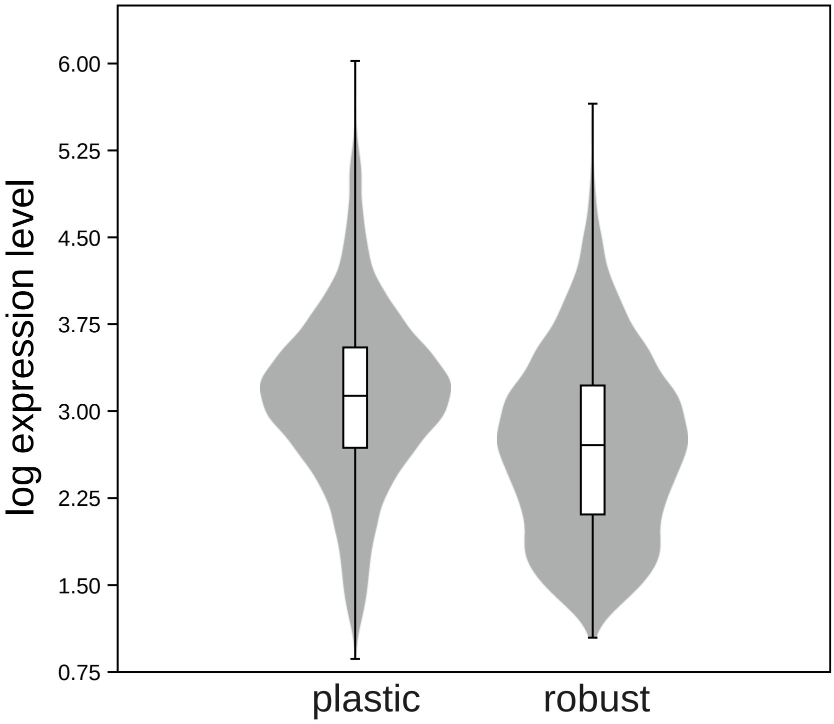

**Supplemental Figure 2.** Relation between pleiotropy proxy and the log mean expression of a gene.

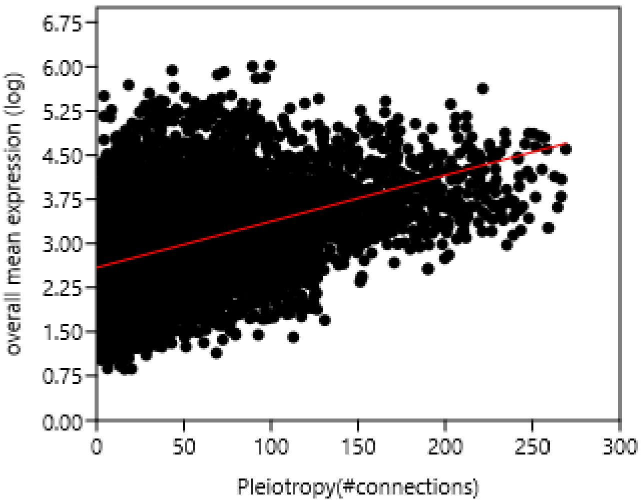

**Supplemental Figure 3**. Bayesian 2-way ANOVA on strength of CGV.

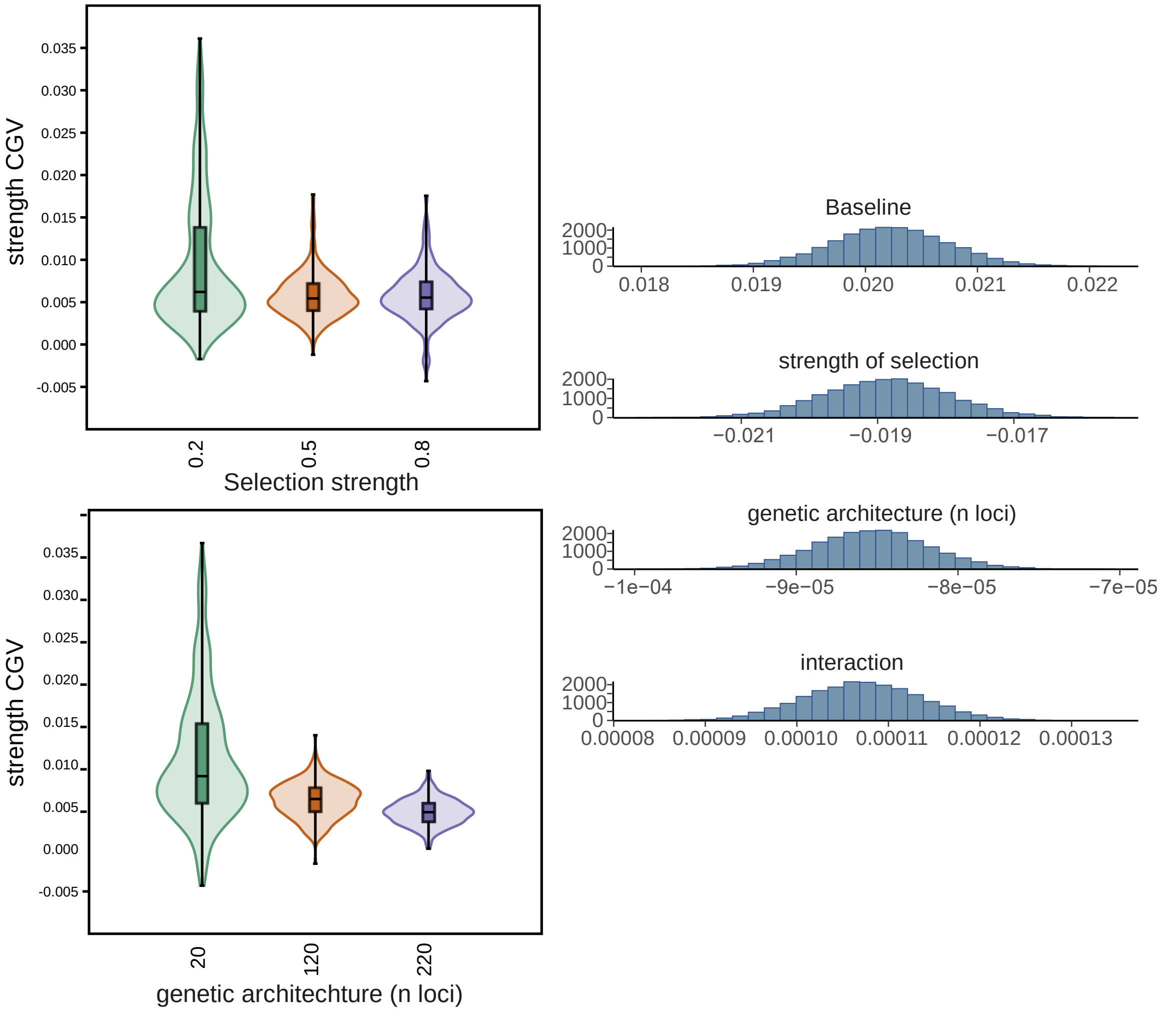

#### Part B Supplemental Tables

**Supplemental Table 1**. Numerical Statistics for Run Lengths per Category

| Category | Mean | Std Dev | 95% CI of Mean |
|---|---|---|---|
| **linear increase** | 1.15 | 0.42 | (1.12, 1.18) |
| **linear decrease** | 1.18 | 0.48 | (1.14, 1.21) |
| **quadratic downwards** | 1.16 | 0.45 | (1.12, 1.19) |
| **quadratic upwards** | 1.20 | 0.50 | (1.17, 1.24) |
| **exponential increase** | 1.25 | 0.58 | (1.22, 1.29) |
| **exponential decrease** | 1.18 | 0.47 | (1.14, 1.21) |
| **MM increase** | 1.32 | 0.64 | (1.28, 1.35) |

**Supplemental Table 2.** Tukey's HSD Post-Hoc Test Results. Multiple Comparison of Means - Tukey HSD, FWER=0.05

| group1 | group2 | meandiff p-adj | lower | upper | reject | |
|---|---|---|---|---|---|---|
| MM increase | exponential decrease | -0.1426 | 0.0 | -0.2122 | -0.073 | True |
| MM increase | exponential increase | -0.0633 | 0.051 | -0.1267 | 0.0001 | False |
| MM increase | linear decrease | -0.1423 | 0.0 | -0.211 | -0.0736 | True |
| MM increase | linear increase | -0.1681 | 0.0 | -0.2358 | -0.1004 | True |
| MM increase | quadratic downwards | -0.1625 | 0.0 | -0.2371 | -0.0879 | True |
| MM increase | quadratic upwards | -0.114 | 0.0 | -0.1814 | 0.0466 | True |
| exponential decrease | exponential increase | 0.0793 | 0.0228 | 0.0064 | 0.1522 | True |
| exponential decrease | linear decrease | 0.0003 | 1.0 | -0.0773 | 0.0778 | False |
| exponential decrease | linear increase | -0.0255 | 0.958 | -0.1022 | 0.0512 | False |
| exponential decrease | quadratic downwards | -0.0199 | 0.9921 | -0.1028 | 0.0629 | False |
| exponential decrease | quadratic upwards | 0.0285 | 0.9278 | -0.0479 | 0.105 | False |
| exponential increase | linear decrease | -0.079 | 0.0209 | -0.1511 | -0.007 | True |
| exponential increase | linear increase | -0.1048 | 0.0003 | -0.1759 | 0.0337 | True |
| exponential increase | quadratic downwards | -0.0992 | 0.0032 | -0.177 | 0.0215 | True |
| exponential increase | quadratic upwards | -0.0508 | 0.3446 | -0.1216 | 0.0201 | False |
| linear decrease | linear increase | -0.0258 | 0.9536 | -0.1016 | 0.0501 | False |
| linear decrease | quadratic downwards | -0.0202 | 0.9911 | -0.1023 | 0.0619 | False |

| | | | | | | |
|---|---|---|---|---|---|---|
| linear decrease | quadratic upwards | 0.0283 | 0.9273 | -0.0473 | 0.1039 | False |
| linear increase | quadratic downwards | 0.0056 | 1.0 | -0.0757 | 0.0868 | False |
| linear increase | quadratic upwards | 0.0541 | 0.3318 | -0.0206 | 0.1287 | False |
| quadratic downwards | quadratic upwards | 0.0485 | 0.5721 | -0.0325 | 0.1295 | False |

**Supplemental Table 3.** Correlations of temperature-wise standard deviations along PCA axes with the temperature deviations from the selective optimum at 16°C.

| | r | p |
|---|---|---|
| **PCA1** | 0.97 | 0.01 |
| **PCA2** | 0.55 | 0.34 |
| **PCA3** | -0.33 | 0.59 |
| **PCA4** | 0.58 | 0.31 |
| **PCA5** | 0.45 | 0.45 |

#### Part C Supplemental Lists

**Supplemental List 1**. Enriched GO-terms among the genes for the different thermal expression reaction norms.

A) Enriched GO terms among genes with a linear positive response to temperature

- cellular nitrogen compound biosynthetic process
- nuclear-transcribed mRNA poly(A) tail shortening
- organic cyclic compound biosynthetic process
- regulation of mRNA metabolic process
- positive regulation of RNA metabolic process
- positive regulation of mRNA catabolic pr...
- aromatic compound biosynthetic process
- tricarboxylic acid metabolic process
- heterocycle biosynthetic process
- positive regulation of nucleobase-contai...
- mitochondrial membrane organization
- organic hydroxy compound metabolic proc...
- triglyceride catabolic process
- amino acid transport
- cytoplasmic translation
- neutral lipid catabolic process
- monatomic cation transport
- DNA-templated transcription

B) Enriched GO terms of genes with linear negative response to temperature.

- negative regulation of biosynthetic proc...
- negative regulation of cellular biosynth...
- negative regulation of metabolic process
- negative regulation of macromolecule met...
- negative regulation of cellular process
- negative regulation of biological proces...
- monatomic ion transmembrane transport
- negative regulation of gene expression
- regulation of carbohydrate metabolic pro...
- negative regulation of macromolecule bio...
- regulation of carbohydrate biosynthetic ...
- monatomic ion transport
- regulation of cellular component size
- actin filament organization
- recombinational repair
- regulation of biosynthetic process
- inorganic ion import across plasma membr...
- inorganic ion transmembrane transport

C) Enriched GO terms of genes with quadratic upward response to temperature.

- neurotransmitter receptor transport to p...
- establishment of protein localization to...
- protein localization to postsynaptic mem...
- organonitrogen compound biosynthetic pro...

- protein localization to cell junction
- neurotransmitter receptor transport
- glutathione catabolic process
- amino acid biosynthetic process
- alpha-amino acid biosynthetic process
- cell junction assembly
- regulation of amide metabolic process
- endoplasmic reticulum unfolded protein r...
- regulation of synapse structure or activ...
- receptor localization to synapse
- positive regulation of transcription elo...
- regulation of postsynaptic membrane neur...
- RNA decapping
- regulation of translation
- carboxylic acid biosynthetic process

D) Enriched GO terms of genes with quadratic downward response to temperature.

- organic substance transport
- RNA splicing, via endonucleolytic cleava...
- intracellular transport
- nitrogen compound transport
- regulation of programmed cell death
- positive regulation of cation transmembr...
- regulation of monatomic ion transmembra...
- positive regulation of transmembrane tra...
- positive regulation of monatomic ion tr...
- negative regulation of programmed cell d...
- regulation of transmembrane transport
- regulation of RNA biosynthetic process
- response to oxygen-containing compound
- homoeostatic process
- regulation of DNA-templated transcriptio...
- positive regulation of RNA biosynthetic ...
- positive regulation of monatomic ion tr... (smaller instance)
- establishment of localization in cell
- intracililary transport

E) Enriched GO terms of genes with exponential positive response to temperature

- positive regulation of macromolecule bio...
- sphingolipid biosynthetic process
- positive regulation of cell cycle
- organic hydroxy compound catabolic proce...
- positive regulation of biosynthetic proc...
- positive regulation of cellular biosynth...
- regulation of cell cycle process
- sphingolipid metabolic process
- positive regulation of transcription by ...
- ceramide biosynthetic process
- positive regulation of cell cycle proces...
- amino sugar catabolic process (larger instance)
- protein modification process

- ceramide metabolic process
- amino sugar metabolic process (smaller instance)
- regulation of chromosome organization
- positive regulation of macromolecule met...
- response to endoplasmic reticulum stress
- modified amino acid transport
- actin nucleation

F) Enriched GO terms of genes with exponential negative response to temperature.

- microtubule polymerization
- retrograde vesicle-mediated transport, G...
- nitrogen compound transport
- regulation of membrane lipid distribu...
- oligopeptide import across plasma membra...
- microtubule polymerization or depolymeri...
- phospholipid transport
- carbohydrate biosynthetic process
- glutathione metabolic process
- mRNA export from nucleus
- RNA localization
- oligopeptide transmembrane transport
- DNA repair
- RNA catabolic process
- macromolecule localization
- mitochondrial translation
- lipid translocation
- organic substance transport
- establishment of RNA localization
- cellular component organization

G) Enriched GO terms of genes with Michealis-Menten-like response to temperature.

- regulation of catalytic activity
- microtubule-based process
- microtubule organizing center organization
- endocytosis
- RNA 3'-end processing
- regulation of MAPK cascade
- DNA recombination
- double-strand break repair
- centriole replication
- regulation of centrosome cycle
- sensory perception of sweet taste

**Supplemental List 2**. Enriched GO-terms in gene sets for CGV & plasticity, CGV & robust, no CGV, & plastic, no CGV & robust.

A) Thermally plastic genes with CGV

- Cellular component assembly
- Cellular component biogenesis
- Protein maturation
- Protein localization to membrane
- Animal organ morphogenesis
- Monocarboxylic acid metabolic process
- Proteolysis involved in protein catabolism
- Establishment of protein localization to...
- Mitochondrial respiratory chain complex...
- Response to hypoxia
- Response to reactive oxygen species
- Protein catabolic process
- Protein acylation
- Biosynthetic process
- Biosynthetic process
- Monatomic cation transport
- Tetrapyrole metabolic process
- Tetrapyrole biosynthetic process
- Positive regulation of phosphorylation
- Regulation of vesicle-mediated transport

B) Genes not thermally plastic with CGV

- alpha-amino acid biosynthetic process
- intrinsic apoptotic signaling pathway
- amino acid biosynthetic process
- regulation of biological quality
- cell adhesion
- cellular response to reactive oxygen spe...
- small molecule biosynthetic process
- proteasome-mediated ubiquitin-dependent ...
- intracellular chemical homeostasis
- aspartate family amino acid metabolic pr...
- proton motive force-driven mitochondrial...
- exonucleolytic catabolism of deadenyl/ate...
- cell projection organization
- apoptotic signaling pathway
- glutathione catabolic process
- regulation of mRNA catabolic process
- meiotic cell cycle
- cellular response to stress
- organonitrogen compound biosynthetic pro...
- rRNA 3'-end processing

C) Thermally plastic without CGV.

- regulation of cell development

- regulation of neurogenesis
- developmental growth
- regulation of extent of cell growth
- defense response to other organism
- positive regulation of intracellular sig...
- neuron projection extension
- synaptic transmission, glutamatergic
- developmental growth involved in morphog...
- axon extension
- developmental cell growth
- regulation of cell differentiation
- response to external stimulus
- synaptic signaling
- regulation of nervous system development
- regulation of response to stimulus
- positive regulation of TOR signaling
- secondary alcohol biosynthetic process
- monatomic anion transport
- tRNA modification

D) Robust without CGV.

- cellular nitrogen compound biosynthetic
- positive regulation of RNA metabolic pro...
- positive regulation of cellular biosynth...
- positive regulation of RNA biosynthetic ...
- positive regulation of macromolecule bio...
- positive regulation of biosynthetic proc...
- positive regulation of nucleobase-contai...
- maturation of LSU-rRNA
- positive regulation of DNA-templated tra...
- organic cyclic compound biosynthetic pro...
- potassium ion import across plasma membr...
- peptide biosynthetic process
- phosphatidyl ethanolamine metabolic proce...
- phosphatidyl ethanolamine biosynthetic pr...
- regulation of cellular biosynthetic proc...
- aromatic compound biosynthetic process
- amino acid catabolic process
- heterocycle biosynthetic process
- positive regulation of nitrogen compound...

### Part D Supplemental Scripts

**Supplemental Script 1**. Python script for fitting different phenotypic response curve functions on gene-expression or other trait data.

```
'''
Python script for fitting different phenotypic response curve functions on gene-expression or
other trait data
Written in Python 3.9
Markus Pfenninger June 2024

'''
### structure of input data:
### csv or tab delimited file
### traits,environmental data (rows) by individuals/samples (columns)
### 1st line: environmental parameter
### 1st column: name of environmental parameter, followed by the environmental parameter for
each individuals/sample
### 2nd line and following: traits
### 1st column: name of the trait, followed by the trait values
### example:
### temp,5,5,5,10,10,....
### size,4,3,3,6,7,6,....

# import necessary modules

import csv
import numpy as np
from scipy.stats import ttest_1samp
from scipy.stats import fisher_exact
from scipy.optimize import curve_fit
from sklearn.metrics import mean_squared_error
from math import log
import matplotlib.pyplot as plt

# create necessary arrays and initiate variables

env = [] #array for the environmental parameter
trait_name = [] #array for the names of the traits
response_func = []
response_func_dir = []
bow_tie = []
positive_slope = 0
negative_slope = 0

def subgroup_statistics(trait_value, env):
    '''
    Function to calculate means and standard deviations for each subgroup defined by the
string array.
    It then compares the subgroups with minimal and maximal means using a one-sided t-test
    to see whether at least two conditions are significantly different.

    Parameters:
    - trait_value: list of float
        List containing float values.
    - env: list of float
        List containing env values to define subgroups.

    Returns:
    - tuple:
        A tuple containing the results of the t-test comparison between subgroups with minimal
and maximal means.
        The tuple format is (t_statistic, p_value).
    '''

    # Creating a dictionary to store subgroups based on string values
    subgroup_dict = {}
    for i in range(len(env)):
        if env[i] not in subgroup_dict:
            subgroup_dict[env[i]] = []
        subgroup_dict[env[i]].append(trait_value[i])

    # Creating field for storage of standard deviations
    stdgroups = []
    cvgroups =[]
```

```
    # Calculating means and standard deviations for each subgroup
    subgroup_stats = {}
    for key, values in subgroup_dict.items():
        subgroup_mean = np.mean(values)
        subgroup_std = np.std(values)
        subgroup_cv = np.std(values)/np.mean(values)
        subgroup_stats[key] = (subgroup_mean, subgroup_std, subgroup_cv)
        stdgroups.append(subgroup_std)
        cvgroups.append(subgroup_cv)

    # Fitting a linear function to the standard deviations per group versus the absolute
distance to optimal environmental condition
    x = [11,6,1,4,9]
    y = stdgroups
    bow_linear, _ = curve_fit(linear_func, x, y)
    y = cvgroups
    stdbowlin, _ = curve_fit(linear_func, x, y)

    # Finding subgroups with minimal and maximal means
    min_mean_group = min(subgroup_stats, key=lambda k: subgroup_stats[k][0])
    max_mean_group = max(subgroup_stats, key=lambda k: subgroup_stats[k][0])

    # Performing one-sided t-test between subgroups with minimal and maximal means
    t_statistic, p_value = ttest_1samp(subgroup_dict[max_mean_group],
np.mean(subgroup_dict[min_mean_group]))

    return t_statistic, p_value, bow_linear, stdbowlin

def linear_func(x, a, b):
    """
    Linear function to fit the data.

    Parameters:
    - x: array-like
        Independent variable.
    - a: float
        Slope of the linear function.
    - b: float
        Intercept of the linear function.

    Returns:
    - array-like:
        Predicted values based on the linear function.
    """
    return a * x + b

def quadratic_func(x, a, b, c):
    """
    Quadratic function to fit the data.

    Parameters:
    - x: array-like
        Independent variable.
    - a: float
        quadratic constant.
    - b: float
        linear constant.
    - c: float
        scaling.

    Returns:
    - array-like:
        Predicted values based on the quadratic function.
    """
    return a * x**2 + b * x + c

def logistic_func(x, a, b, c):
    """
    Logistic function to fit the data.

    Parameters:
    - x: array-like
        Independent variable.
    - a: float
        max of function.
    - b: float
        slope.
    - c: float
```

```
        midpoint.

    Returns:
    - array-like:
        Predicted values based on the logistic function.
    """
    return a / (1 + b * np.exp(x * c))

def MichMent_func(x, a, b):
    """
    Michaelis-Menten function to fit the data.

    Parameters:
    - x: array-like
        Independent variable.
    - a: float
        max of function.
    - b: float
        slope.

    Returns:
    - array-like:
        Predicted values based on the Michaelis-Menten function.
    """
    return (a * x) / (b + x)

def exponential_func(x, a, b, c):
    """
    Exponential function to fit the data.

    Parameters:
    - x: array-like
        Independent variable.
    - a: float
        Coefficient of the exponential function.
    - b: float
        Exponential base.

    Returns:
    - array-like:
        Predicted values based on the exponential function.
    """
    return a * np.exp(b * x) + c

def beta_func(x, a, b, c):
    """
    Probability density function of the beta distribution: f(x) = c * x**(a-1) * (x-1)^(b-1)
    constrained on a being larger than b

    Parameters:
    - x: float
        The input value.
    - a: float
        initial increase
    - b: float
        decline rate
    - c: float
        constant

    Returns:
    - float:
        The output value of the beta function at x.
    """
    return c * x**(a-1) * (x-1)**(b-1)

def fit_and_compare(x, y):
    """
    Fits a linear and an exponential function to the data, compares the fits using AIC, and
returns the parameters of the better fit.

    Parameters:
    - x: array-like
        Independent variable.
    - y: array-like
        Dependent variable.
    - y_true: array-like
        True values for comparison.
```

```
    Returns:
    - tuple:
        Parameters of the better fit
    - chosen function
    - AIC.
    """
    # Array for AICs
    model_select = []
    model_params = []
    model_names = ["linear", "quadratic", "logistic","Michaelis-Menten","exponential","beta"]

    # Fit linear function
    popt_linear, _ = curve_fit(linear_func, x, y)
    residuals_linear = y - linear_func(x, *popt_linear)
    linear_aic = len(y) * np.log(np.sum(residuals_linear**2) / len(y)) + 2 * 2  # AIC for
linear model
    model_select.append(linear_aic)
    model_params.append(popt_linear)

    # Fit quadratic function
    popt_quadratic, _ = curve_fit(quadratic_func, x, y)
    residuals_quadratic = y - quadratic_func(x, *popt_quadratic)
    quadratic_aic = len(y) * np.log(np.sum(residuals_quadratic**2) / len(y)) + 2 * 3  # AIC
for linear model
    model_select.append(quadratic_aic)
    model_params.append(popt_quadratic)

    # Fit logistic function
    popt_logistic, _ = curve_fit(logistic_func, x, y, bounds=(0,35))
    residuals_logistic = y - logistic_func(x, *popt_logistic)
    logistic_aic = len(y) * np.log(np.sum(residuals_logistic**2) / len(y)) + 2 * 3  # AIC for
linear model
    model_select.append(logistic_aic)
    model_params.append(popt_logistic)

    # Fit MichMent function
    popt_MichMent, _ = curve_fit(MichMent_func, x, y)
    residuals_MichMent = y - MichMent_func(x, *popt_MichMent)
    MichMent_aic = len(y) * np.log(np.sum(residuals_MichMent**2) / len(y)) + 2 * 2  # AIC for
linear model
    model_select.append(MichMent_aic)
    model_params.append(popt_MichMent)

    # Fit exponential function
    try:
       popt_expo, _ = curve_fit(exponential_func, x, y, maxfev = 5000)
    except RuntimeError:
       model_select.append(100000)
       model_params.append("Nan")
    else:
       residuals_expo = y - exponential_func(x, *popt_expo)
       expo_aic = len(y) * np.log(np.sum(residuals_expo**2) / len(y)) + 2 * 2  # AIC for
linear model
       model_select.append(expo_aic)
       model_params.append(popt_expo)

    # Fit beta function
    popt_beta, _ = curve_fit(beta_func, x, y, bounds=(0,35))
    if popt_beta[0] > popt_beta[1]:
        residuals_beta = y - beta_func(x, *popt_beta)
        beta_aic = len(y) * np.log(np.sum(residuals_beta**2) / len(y)) + 2 * 3  # AIC for beta
model
        model_select.append(beta_aic)
        model_params.append(popt_beta)
    else:
        model_select.append(10e+100)
        model_params.append("N.a")

    #Compare AICs
    best_AIC = min(model_select)
    ind = model_select.index(best_AIC)
    best_model = model_names[ind]
    parameters = model_params[ind]
    return best_model, best_AIC, parameters

def plot_pie_chart(response_func, file_name):
    """
```

```
    Function to plot a labeled pie chart for the frequency of items in a response_func of
strings.

    Parameters:
    - response_func: list
        A list of strings representing the items for which the frequency needs to be
visualized.

    Returns:
    - None
        This function doesn't return anything but displays the pie chart.

    Raises:
    - ValueError:
        Raises an error if the input response_func is empty.
    """

    # Checking if the input response_func is empty
    if not response_func:
        raise ValueError("Input response_func is empty. Please provide a non-empty list of
strings.")

    # Counting the frequency of each unique item in the response_func
    frequency_dict = {}
    for item in response_func:
        if item in frequency_dict:
            frequency_dict[item] += 1
        else:
            frequency_dict[item] = 1

    # Extracting labels and sizes for the pie chart
    labels = frequency_dict.keys()
    sizes = frequency_dict.values()

    # Plotting the pie chart
    plt.figure(figsize=(8, 8))
    plt.pie(sizes, labels=labels, autopct='%1.1f%%', startangle=140)
    plt.axis('equal')  # Equal aspect ratio ensures that pie is drawn as a circle.

    # Adding title to the pie chart
    plt.title('Frequency of different reaction norm functions')

    # Displaying and saving the pie chart
    plt.savefig(str(file_name) + ".pdf", format="pdf", bbox_inches="tight")
    plt.show()

def response_direction(response, best_fit_params):
    """
    This function splits the general response functions further into the direction of response

    Args:
    -the inferred response
    - a tuple with the best-fit parameters

        Raises:
    - ValueError:
        Raises an error if the input response_func is empty.
    """

    # Checking if the input response_func is empty
    if not response_func:
        raise ValueError("Input response_func is empty. Please provide a non-empty list of
strings.")

    # checking the response function and inferring the direction from the parameters
    ["linear", "quadratic", "logistic","Michaelis-Menten","exponential","beta"]

    if response == "linear":
       if best_fit_params[2][0] > 0:
              response_func_dir.append("linear increase")
              res_dir = "linear increase"
       else:
              response_func_dir.append("linear decrease")
              res_dir = "linear decrease"

    elif response == "quadratic":
       if best_fit_params[2][0] > 0:
              response_func_dir.append("quadratic upwards")
```

```
                res_dir = "quadratic upwards"
        else:
                response_func_dir.append("quadratic downwards")
                res_dir = "quadratic downwards"

    elif response == "logistic":
        if best_fit_params[2][0] > 0 and best_fit_params[2][2] > 0:
                response_func_dir.append("logistic down")
                res_dir = "logistic down"
        elif best_fit_params[2][0] > 0 and best_fit_params[2][2] < 0:
                response_func_dir.append("logistic up")
                res_dir = "logistic up"
        elif best_fit_params[2][0] < 0 and best_fit_params[2][2] > 0:
                response_func_dir.append("logistic up")
                res_dir = "logistic up"
        elif best_fit_params[2][0] < 0 and best_fit_params[2][2] < 0:
                response_func_dir.append("logistic down")
                res_dir = "logistic down"

    elif response == "Michaelis-Menten":
         if best_fit_params[2][0] > 0:
                response_func_dir.append("MM increase")
                res_dir = "MM increase"
         else:
                response_func_dir.append("MM downwards")
                res_dir = "MM downwards"

    elif response == "exponential":
                if best_fit_params[2][0] > 0 and best_fit_params[2][1] > 0:
                        response_func_dir.append("exponential increase")
                        res_dir = "exponential increase"
                elif best_fit_params[2][0] < 0 and best_fit_params[2][1] < 0:
                        response_func_dir.append("exponential increase")
                        res_dir = "exponential increase"
                else:
                        response_func_dir.append("exponential decrease")
                        res_dir = "exponential decrease"
    elif response == "beta":
        response_func_dir.append("beta")
        res_dir = "beta"
    else:
        res_dir = "Nan"
    return(res_dir)

def plot_linear_functions(functions):
  """
  This function takes an array of tuples, representing the parameters of linear functions,
  and plots them for a range of 0-35 in a single plot.

  Args:
    functions: An array of tuples, where each tuple represents the parameters (slope,
intercept)
               of a linear function.
  """
  x = np.linspace(0, 20, 100)  # Create 100 points between 0 and 35

  for slope, intercept in functions:
    y = slope * x + intercept  # Calculate y values for each function
    plt.plot(x, y, label=f"y = {slope}x + {intercept}")  # Plot the function with a label

  plt.xlabel("Deviation from optimal temperature")
  plt.ylabel("Standardised interindividual transcription variability")
  plt.savefig("Bowtie.pdf", format="pdf", bbox_inches="tight")
  plt.show()

def fisher_exact_test(x, y):
  """
  This function performs Fisher's exact test on two contingency tables represented by x and y.

  Args:
    x: An integer representing the number of successes in the first table.
    y: An integer representing the number of successes in the second table.

  Returns:
    A tuple containing the p-value and odds ratio of the Fisher's exact test.
  """
  # Assuming the tables have equal counts of failures (e.g., 2x2 tables)
  table = [[x, y], [(x + y)/2, (x + y)/2]]  # Create the contingency table
```

```
  odds_ratio, p_value = fisher_exact(table)
  return p_value, odds_ratio

### Main
# read in input file and prepare the output file

infile = "in.txt"            #input file, structured as detailed above

outfname = "All_genes_results.out"
outf = open(outfname, "w")
outf.write("geneID\tbowtie_slope\tbowtie_intercept\tresponse\tdirection\parameters\n")


with open(infile) as to_read:
        reader = csv.reader(to_read, delimiter = "\t")
        #get the environmental parameter and write in array
        header = next(reader)
        for i in range(1,len(header)):
                env.append(float(header[i]))
        env = np.array(env)
        for row in reader:
                trait_name.append(row[0])
                outf.write(str(row[0]) + "\t")
                trait = []
                for i in range(1,len(row)):
                        trait.append(float(row[i]))
                trait = np.array(trait)

                # Calculating statistics and performing t-test
                t_stat, p_val, bow_linear, stdbowlin = subgroup_statistics(trait, env)

                # Adding parameters of s.d. vs. deviation from optimum to array  and write them
to file
                bow_tie.append(bow_linear)
                outf.write(str(bow_linear[0]) + "\t" + str(bow_linear[1]) + "\t" +
str(stdbowlin[0]) + "\t")
                if bow_linear[0] > 0:
                        positive_slope = positive_slope + 1
                elif bow_linear[0] < 0:
                        negative_slope = negative_slope + 1

                # test, if at least two conditions are significantly different
                if p_val <= 0.05:
                        # calling function fitting function
                        best_fit_params = fit_and_compare(env, trait)
                        print(row[0], " ", best_fit_params[0], " ", best_fit_params[1], " ",
best_fit_params[2])
                        response = best_fit_params[0]
                elif p_val > 0.05:
                        print(row[0]," no difference among environments")
                        response = "flat"
                response_func.append(response)
                # calling fine split function
                direction = response_direction(response, best_fit_params)
                outf.write(str(response) + "\t" + str(direction) + "\t")
                for i in range(0, len(best_fit_params[2])):
                        outf.write(str(best_fit_params[2][i]) + "\t")
                outf.write("\n")

# calling Fisher's exact test
p_value, odds_ratio = fisher_exact_test(positive_slope, negative_slope)
print("number of positive slopes: ", positive_slope, "number of negative slopes: ",
negative_slope)
print(p_value, "  ", odds_ratio)
outf.write("positive slopes \t negative slopes \n")
outf.write(str(positive_slope) + "\t" + str(negative_slope) + "\n")
outf.write("p value: \t" + str(p_value) + "\n")
outf.write("odds ratio \t" + str(odds_ratio) + "\n")

# calling plotting functions
plot_linear_functions(stdbowlin)
file_name = "response_function"
plot_pie_chart(response_func, file_name)
file_name = "response_function_direction"
plot_pie_chart(response_func_dir, file_name)
outf.close()
```

**Supplemental script 2**. Runs-test with subsequent ANOVA and Tukey's post-hoc test.

```
import random
import matplotlib.pyplot as plt
import numpy as np
import seaborn as sns # For enhanced plotting
import pandas as pd   # For easier data manipulation for plotting
import scipy.stats as stats # For statistical functions, especially confidence intervals
from statsmodels.stats.multicomp import pairwise_tukeyhsd # For post-hoc test

def count_runs(sequence):
    """
    Counts the total number of runs in a given sequence.
    A run is a consecutive series of identical elements.
    """
    if not sequence:
        return 0

    runs = 1
    for i in range(1, len(sequence)):
        if sequence[i] != sequence[i-1]:
            runs += 1
    return runs

def get_run_lengths_by_category(sequence):
    """
    Records the lengths of runs for each category in a sequence.

    Args:
        sequence (list): The sequence of categories.

    Returns:
        dict: A dictionary where keys are categories and values are lists
              of run lengths for that category.
              Example: {'A': [3, 1], 'B': [2], 'C': [4, 1]}
    """
    run_lengths_by_category = {}
    if not sequence:
        return run_lengths_by_category

    current_category = sequence[0]
    current_run_length = 1

    for i in range(1, len(sequence)):
        if sequence[i] == current_category:
            current_run_length += 1
        else:
            # End of a run, record it
            if current_category not in run_lengths_by_category:
                run_lengths_by_category[current_category] = []
            run_lengths_by_category[current_category].append(current_run_length)

            # Start a new run
            current_category = sequence[i]
            current_run_length = 1

    # After the loop, record the last run
    if current_category not in run_lengths_by_category:
        run_lengths_by_category[current_category] = []
    run_lengths_by_category[current_category].append(current_run_length)

    return run_lengths_by_category

def permutation_runs_test(observed_sequence, n_permutations=10000):
    """
    Performs a permutation-based runs test for a sequence with multiple categories.

    Args:
        observed_sequence (list): The observed sequence of categories (e.g., ['A', 'A', 'B',
'C', ...]).
        n_permutations (int): The number of permutations to perform.
```

```
    Returns:
        tuple: A tuple containing:
            - observed_runs (int): The number of runs in the observed sequence.
            - p_value_clustering (float): P-value for the hypothesis of clustering (fewer
runs).
            - p_value_alternation (float): P-value for the hypothesis of alternation (more
runs).
            - permuted_runs_counts (list): A list of run counts from all permutations.
    """
    observed_runs = count_runs(observed_sequence)
    print(f"Observed total number of runs: {observed_runs}")

    permuted_runs_counts = []

    for _ in range(n_permutations):
        shuffled_sequence = list(observed_sequence)
        random.shuffle(shuffled_sequence)

        permuted_runs_counts.append(count_runs(shuffled_sequence))

    p_value_clustering = sum(1 for runs in permuted_runs_counts if runs <= observed_runs) /
n_permutations
    p_value_alternation = sum(1 for runs in permuted_runs_counts if runs >= observed_runs) /
n_permutations

    print(f"P-value for clustering (fewer total runs): {p_value_clustering:.4f}")
    print(f"P-value for alternation (more total runs): {p_value_alternation:.4f}")

    return observed_runs, p_value_clustering, p_value_alternation, permuted_runs_counts

# --- Example Usage ---
if __name__ == "__main__":
    # Example genomic feature sequence with 8 categories (A-H)
    # Let's make one category (e.g., 'A') clearly clustered to see the effect
    my_genomic_sequence = [    ]
    # You would replace this with your actual data

    print("--- Running Permutation Test for Total Runs ---")
    obs_runs, p_cluster, p_alternation, perm_counts = permutation_runs_test(
        my_genomic_sequence, n_permutations=20000
    )

    # --- Visualization of Total Runs Distribution ---
    plt.figure(figsize=(10, 6))
    plt.hist(perm_counts, bins=30, color='skyblue', edgecolor='black', alpha=0.7)
    plt.axvline(obs_runs, color='red', linestyle='dashed', linewidth=2, label=f'Observed Runs:
{obs_runs}')
    plt.title('Distribution of Total Runs under Randomness (Permutation Test)')
    plt.xlabel('Number of Runs')
    plt.ylabel('Frequency')
    plt.legend()
    plt.grid(axis='y', alpha=0.75)
    plt.show()

    # --- Interpretation of Overall Runs Test ---
    alpha = 0.05
    print("\n--- Interpretation of Overall Runs Test ---")
    if p_cluster < alpha:
        print(f"P-value for clustering ({p_cluster:.4f}) is less than alpha ({alpha}).")
        print("This suggests that the features are significantly clustered overall (fewer
total runs than expected by chance).")
    elif p_alternation < alpha:
        print(f"P-value for alternation ({p_alternation:.4f}) is less than alpha ({alpha}).")
        print("This suggests that the features show a significant alternating pattern overall
(more total runs than expected by chance).")
    else:
        print(f"Neither clustering (p={p_cluster:.4f}) nor alternation (p={p_alternation:.4f})
is statistically significant at alpha={alpha}.")
        print("The observed sequence of features appears random with respect to total runs.")


    print("\n--- Analyzing Run Length Distribution per Category ---")
    # Get run lengths for each category in the observed sequence
```

```
    category_run_lengths = get_run_lengths_by_category(my_genomic_sequence)

    # Convert the dictionary to a pandas DataFrame for easy plotting with seaborn
    data_for_plot = []
    for category, lengths in category_run_lengths.items():
        for length in lengths:
            data_for_plot.append({'Category': category, 'Run Length': length})

    df_run_lengths = pd.DataFrame(data_for_plot)

    if not df_run_lengths.empty:
        # Sort categories for consistent plotting order (optional)
        # sorted_categories = sorted(df_run_lengths['Category'].unique())
        # df_run_lengths['Category'] = pd.Categorical(df_run_lengths['Category'],
categories=sorted_categories, ordered=True)

        plt.figure(figsize=(12, 7))
        sns.boxplot(x='Category', y='Run Length', data=df_run_lengths, palette='viridis',
showfliers=False)
        sns.stripplot(x='Category', y='Run Length', data=df_run_lengths, color='black',
size=4, jitter=0.2, alpha=0.6)

        plt.title('Run Length Distribution per Category in Observed Sequence')
        plt.xlabel('Genomic Feature Category')
        plt.ylabel('Run Length')
        plt.grid(axis='y', linestyle='--', alpha=0.7)
        plt.xticks(rotation=45, ha='right')
        plt.tight_layout()
        plt.show()

        print("\n--- Numerical Statistics for Run Lengths per Category ---")
        print("{:<15} {:<10} {:<10} {:<25}".format("Category", "Mean", "Std Dev", "95% CI
(Mean)"))
        print("-" * 60)

        # List to store data for ANOVA
        anova_data = []
        # List to store groups for ANOVA (each list of run lengths)
        anova_groups = []

        for category, lengths in category_run_lengths.items():
            if lengths: # Ensure there are run lengths for the category
                np_lengths = np.array(lengths)
                mean_len = np.mean(np_lengths)
                std_dev_len = np.std(np_lengths, ddof=1) # ddof=1 for sample standard
deviation

                n = len(np_lengths)
                if n > 1:
                    std_err_mean = std_dev_len / np.sqrt(n)
                    ci_lower, ci_upper = stats.t.interval(0.95, df=n-1, loc=mean_len,
scale=std_err_mean)
                    ci_str = f"({ci_lower:.2f}, {ci_upper:.2f})"
                else:
                    std_dev_len = float('nan')
                    ci_str = "N/A (single run)"

                print(f"{category:<15} {mean_len:<10.2f} {std_dev_len:<10.2f} {ci_str:<25}")

                # Prepare data for ANOVA: only include categories with at least 1 run (already
handled by 'if lengths')
                anova_groups.append(np_lengths)

            else:
                print(f"{category:<15} {'N/A':<10} {'N/A':<10} {'N/A':<25}")

        print("\n--- Statistical Tests for Differences in Mean Run Lengths ---")

        # Perform ANOVA if there are at least two groups with data
        if len(anova_groups) >= 2:
            # Filter out groups with insufficient data for ANOVA (e.g., if a category had no
runs)
```

```
            # f_oneway handles groups of size 1, but it's good practice to ensure meaningful
comparisons
            valid_anova_groups = [group for group in anova_groups if len(group) > 0] # Ensure
groups are not empty

            if len(valid_anova_groups) >= 2:
                f_statistic, p_value_anova = stats.f_oneway(*valid_anova_groups)
                print(f"\n--- One-Way ANOVA Test ---")
                print(f"F-statistic: {f_statistic:.4f}")
                print(f"P-value: {p_value_anova:.4e}") # Use scientific notation for small p-
values

                if p_value_anova < alpha:
                    print(f"The ANOVA p-value ({p_value_anova:.4e}) is less than alpha
({alpha}).")
                    print("This indicates that there is a statistically significant difference
in mean run lengths among at least two of the categories.")
                    print("Performing Tukey's HSD post-hoc test to find specific
differences.")

                    # Perform Tukey's HSD post-hoc test
                    # This requires reshaping data into a single column of values and a single
column of groups

                    # Ensure df_run_lengths is not empty before proceeding
                    if not df_run_lengths.empty:
                        try:
                            # pairwise_tukeyhsd handles multiple comparisons
                            tukey_results = pairwise_tukeyhsd(endog=df_run_lengths['Run
Length'],
                                                              groups=df_run_lengths['Category'],
                                                              alpha=alpha)
                            print("\n--- Tukey's HSD Post-Hoc Test Results ---")
                            print(tukey_results)
                            print("\nInterpretation of Tukey's HSD:")
                            print("- 'reject=True' indicates a statistically significant
difference (at alpha={}) between the two categories.".format(alpha))
                            print("- This helps identify which specific categories contribute
to the overall significant ANOVA result.")
                        except ValueError as e:
                            print(f"\nCould not perform Tukey's HSD: {e}")
                            print("This might happen if there are too few data points for some
categories or if all groups have identical means.")
                    else:
                        print("\nCannot perform Tukey's HSD as DataFrame for plotting is
empty.")
                else:
                    print(f"The ANOVA p-value ({p_value_anova:.4e}) is greater than or equal
to alpha ({alpha}).")
                    print("There is no statistically significant difference in mean run
lengths among the categories.")
                    print("Therefore, a post-hoc test is not necessary.")
            else:
                print("\nANOVA requires at least two categories with data for comparison.")
        else:
            print("\nNot enough categories with run length data (need at least 2) to perform
ANOVA.")

        print("\n--- Final Inference Guidance ---")
        print("1. **Overall Run Test:** Look at the first set of p-values to see if the
overall sequence is significantly clustered or dispersed.")
        print("2. **Numerical Statistics (Mean, Std Dev, CI):** Use these to get a
quantitative sense of run lengths for each category. Categories with higher means and CIs that
don't overlap with others are good candidates for driving clustering.")
        print("3. **Boxplots with Jitter:** Visual confirmation of the numerical statistics,
showing the spread and distribution.")
        print("4. **ANOVA (and Tukey's HSD if ANOVA is significant):**")
        print("   - ANOVA's p-value tells you if *any* categories' mean run lengths are
significantly different.")
        print("   - If ANOVA is significant, Tukey's HSD pinpoints *which specific pairs* of
categories have statistically different mean run lengths. This is the definitive answer for
identifying categories that over-proportionally contribute to the pattern (e.g., having
significantly longer runs).")
```

```
    else:
        print("No run lengths to plot or calculate statistics. The sequence might be empty or contains only one run.")
```

**Supplemental Script 3.** Simulation of CGV for a polygenic trait under the assumption that each allele at the contributing loci has its own allelic response curve.

```
##################################################################################
# dynamicCGV Simulation                                                          #
# Markus Pfenninger, Feb 25                                                      #
# written in python3.9                                                           #
##################################################################################

### Module import
from random import randint, random, gauss, uniform, shuffle, sample
import numpy as np
import math
from math import sqrt, pi, exp, sin
from scipy.stats import linregress, fisher_exact, beta
import copy
import csv
import matplotlib.pyplot as plt


###INPUT/OUTPUT MODULE
##Variables
N = 1000                 #adult population size (>larger values, less drift)
x_juv = 10             #factor by which the offspring generation is larger than the adult
population = the clutch size per parent (>larger values, less drift, faster adaptation)
selstr = 0.8   #strength of selection: exponential decline of survivial probability with
deviation from optimal phenotype (>larger values less survival probability with increasing
distance from optimal phenotype)
nloc = 220              #number of loci to contribute to the phenotypic trait (qtl)
generations = 30               #number of generations to run each population (>increases drift
differences)
plot = "y"
env_range = nloc*2
sel_opt = nloc/2
replicates = 100

##creation of arrays for the storage of results
phenopopmean = []
phenopopstd = []
heritability = []
popPheno = [[] for _ in range(N)]
allelic_resp_func = [[] for _ in range(nloc)]



###DEFINED FUNCTIONS

def create_allelic_response_curves(nloc, env_range, allelic_resp_func):
        #for each locus
        for i in range(0,nloc):
                #i) draw a random intercept between -1 and 1
                intercept_A = uniform(0,1)
                #ii) calculate the limits for the max positive and min negative slope for not
to exceed the limits 0 and 1 for #the allelic contributions in the given environmental range 0
- X as
                #- max pos slope = (intercept * -1/X) + 1/X
                #- min neg slope = (intercept * -1/X)
                #iii) draw random a random slope between these two limits
                slope_A = uniform((intercept_A * (-1/env_range)),(intercept_A * (-1/env_range)
+ 1/env_range))
                #iv) for the alternate allele, change intercept and slope by drawing random
values from a gaussian distribution with the old values as means and 0.1 as s.d.
                intercept_a = gauss(intercept_A,0.1)
                slope_a = uniform((intercept_a * (-1/env_range)),(intercept_a * (-1/env_range)
+ 1/env_range))
                allelic_resp_func[i].append(slope_A)
                allelic_resp_func[i].append(intercept_A)
                allelic_resp_func[i].append(slope_a)
                allelic_resp_func[i].append(intercept_a)


        #for i in range(0,len(allelic_resp_func)):
        #       # Plot the estimated line
        #       y_A = []
        #       y_a = []
        #       x = list(range(0,env_range))
        #       for j in range(0,len(x)):
```

```
        #              y_A.append((allelic_resp_func[i][0] * x[j]) + allelic_resp_func[i][1])
        #              y_a.append((allelic_resp_func[i][2] * x[j]) + allelic_resp_func[i][3])
        #       plt.plot(x, y_A, label = "locus 1" + str(i) + " allele A")
        #       plt.plot(x, y_a, label = "locus 1" + str(i) + " allele a")
        #plt.xlabel('enviromental condition')
        #plt.ylabel('allelic contribution to trait')
        #plt.show()
        return (allelic_resp_func)

def initial_population_creation(N, nloc, allelefreq, arf):
        """
        Creates an initial population
        Parameters:
        - N: int
                Size of adult population.
        - nloc: int
                Number of qtl
        -neutl: int
                Number of neutral loci

        Returns:
        - population:
                A list of lists, containing the genotypes of the individuals in the population

            """
        population = [[] for _ in range(N)] #meta field where all the information for an
individual is stored in a subfield
        #creating the initial juvenile population
        for ind in range (0,(N)):
                #assigning a random identifier to the individual (for later randomisation
purposes)
                randID = randint(0,10*N)
                population[ind].append(randID)
                #assign phenotypically relevant alleles to individual
                for loc in range(0,nloc):
                        allele = allelefreq[loc][0]
                        rand = randint(1,100)
                        genotype = 0
                        if rand <= (100 * allele):
                                genotype = genotype + 0
                        else:
                                genotype = genotype + 1
                        rand = randint(1,100)
                        if rand <= (100 * allele):
                                genotype = genotype + 0
                        else:
                                genotype = genotype + 1
                        population[ind].append(genotype)
        #calculate the selectively relevant phenotypic value for the current individual
                phenotype_value = 0
                for loc in range(0,nloc):
                        y = float(population[ind][loc+1])
                        #print(y)
                        if y == 0:
                                phenotype_value = phenotype_value + ((2 * arf[loc][0] * sel_opt
+ arf[loc][1]))
                        elif y == 1:
                                phenotype_value = phenotype_value + ((arf[loc][0] * sel_opt +
arf[loc][1]) + (arf[loc][2] * sel_opt + arf[loc][3]))
                        elif y == 2:
                                phenotype_value = phenotype_value + ((2 * arf[loc][2] * sel_opt
+ arf[loc][3]))
        #and record resulting value
                population[ind].append(phenotype_value)
                #print(phenotype_value)

        #print("size:", len(population))
        return population

def random_mating_selection (population, N, x_juv, nloc, selstr, gen, sel_opt, arf):
        """
        Reproduces and selects the population
        Parameters:
        - N: int
                Size of adult population.
        - x_juv: int
                Number of offspring per adults
        - nloc: int
```

```
		Number of qtl
	-neutl: int
		Number of neutral loci
	Returns:
	- population:
		A list of lists, containing the genotypes and the phenotypic value of the individuals in the population
	"""
	#prepare the initial population for reproduction by copying it to a different array
	adults = copy.deepcopy(population)
	generations = 0
	fam = [[] for _ in range(int(len(adults)/2))]
	for generations in range(0,gen):
		###mate the individuals randomly
		pair = 0
		no_pairs = int(len(adults)/2)
		#empty the current population (i.e. make space for the next generation)
		population = [[] for _ in range(N*x_juv)]
		#pick randomly two individuals from the adults for mating
		for pairs in range(0,no_pairs):
			rand1 = randint(0,len(adults)-1)
			indiv1 = adults[rand1]
			rand2 = randint(0,len(adults)-1)
			indiv2 = adults[rand2]
			if generations == gen-1:
				fam[pairs].append((adults[rand1][nloc+1] + adults[rand2][nloc+1]) / 2)
			#now the pair produces 2*x_juv offspring
			for ind in range (2* x_juv * pairs, 2* x_juv * pairs + 2 * x_juv):
				randID = randint(0,10*N*x_juv)
				population[ind].append(randID)
				#qtl and ntl
				for loc in range(0,nloc):
					#random draw of gametes at each locus
					genotype = 0
					if indiv1[loc + 1] == 0:
						genotype = genotype + 0
					elif indiv1[loc + 1] == 1:
						coin = randint(0,1)
						genotype = genotype + coin
					else:
						genotype = genotype + 1

					if indiv2[loc + 1] == 0:
						genotype = genotype + 0
					elif indiv2[loc + 1] == 1:
						coin = randint(0,1)
						genotype = genotype + coin
					else:
						genotype = genotype + 1

					population[ind].append(genotype)
				#calculate the phenotypic value for the current individual
				phenotype_value = 0
				for loc in range(0,nloc):
					y = float(population[ind][loc+1])
					#print(y)
					if y == 0:
						phenotype_value = phenotype_value + (2 * arf[loc][0] * sel_opt + arf[loc][1])
					elif y == 1:
						phenotype_value = phenotype_value + (arf[loc][0] * sel_opt + arf[loc][1]) + (arf[loc][2] * sel_opt + arf[loc][3])
					elif y == 2:
						phenotype_value = phenotype_value + (2 * arf[loc][2] * sel_opt + arf[loc][3])
				#and record resulting value
				population[ind].append(phenotype_value)
			pair = pair + 1 #reproduction finished, next pair, please
		#empty adult population
		adults = []
		#exponential decrease of probability to survive with distance to selective optimum (= hard selection)
		shuffle(population) #mix population (to avoid that particular families are overrepresented)
		ind = 0
```

```
		try: #under certain parameter combinations, less than N individuals may pass
the selection triage and therefore an index error occurs. Since the purpose of the simulation
is not to infer selection processes, the error is catched
			while len(adults) < N:
				y = random()
				abs_dev = (math.sqrt(np.square(sel_opt -
population[ind][nloc+1])))
				if y < exp(-abs_dev * selstr):
					adults.append(population[ind])
				ind = ind + 1
		except IndexError: #by adding individuals with less stringent selection to the
adult population
			to_add = N - len(adults)
			print("only ", len(adults), " individuals survived selection")
			print(N - len(adults), " individuals are added from the juv. population
of ", len(population))
			while len(adults) < N:
				ind = randint(0,len(population)-1)
				adults.append(population[ind])
		generations = generations + 1
		adults_mean = np.mean(adults, axis=0)
		adults_std = np.std(adults, axis=0)
		#print("generation: ", generations, "mean: ", adults_mean[nloc+1], " s.d.: ",
adults_std[nloc+1])
	return adults

####MAIN

with open("nloc220_08.out", 'w') as outf:
	for reps in range(0,replicates):

		#####GENETIC ARCHITECTURE MODULE
		#all samples start out from the same allele frequencies
		#creation of new population
		#first, the genetic architecture
		allelefreq = [[] for _ in range(nloc)]
		#draw random allele frequency from beta distribution between 0.1 and 0.9 for
each of the loci
		loc = 0
		while loc < nloc:
			freq = beta.rvs(0.5,0.5)
			if freq > 0.10 and freq < 0.90:
				allelefreq[loc].append(freq)
				loc = loc + 1

		arf = create_allelic_response_curves(nloc, env_range, allelic_resp_func)
		#####CREATING THE POPULATION
		#####INITIAL POPULATION CREATION
		#call initial population creation function
		population = initial_population_creation(N, nloc, allelefreq, arf)
		#determining the phenotypic value population parameter values
		initial_mean = np.mean(population, axis=0)
		initial_std = np.std(population, axis=0)
		#print("initial: ",initial_mean[nloc+1], " ", initial_std[nloc+1])
		sel_opt = round(gauss(initial_mean[nloc+1], 1.5))
		#print("selective optimum: ", sel_opt)


		###### REPRODUCTION MODULE
		adults = random_mating_selection (population, N, x_juv, nloc, selstr,
generations, sel_opt, arf)
		#and take a random sample from them
		pop_sample = sample(adults, 30)

		####BOWTIE CAlCULATIONS
		pop_sample_mean = np.mean(pop_sample, axis=0)
		pop_sample_std = np.std(pop_sample, axis=0)
		#print("mean at sel_opt: ", pop_sample_mean[nloc+1], " s.d.: ",
pop_sample_std[nloc+1])

		#field of size individuals (ind) * environmental conditions (cond)
		cgv = [[] for _ in range(len(pop_sample))]
		#loop over all individuals
		for ind in range(0,len(pop_sample)):
			#in all conditions
			for cond in range(0,env_range):
				#and calculate the phenotype value for this condition
				phenotype_value = 0
```

```
                              for loc in range(0,nloc):
                                     y = float(pop_sample[ind][loc+1])
                                     #print(y)
                                     if y == 0:
                                            phenotype_value = phenotype_value + (2 * arf[loc][0] * cond + arf[loc][1])
                                     elif y == 1:
                                            phenotype_value = phenotype_value + (arf[loc][0] * cond + arf[loc][1]) + (arf[loc][2] * cond + arf[loc][3])
                                     elif y == 2:
                                            phenotype_value = phenotype_value + (2 * arf[loc][2] * cond + arf[loc][3])
                              #if cond == sel_opt:
                                     #print(phenotype_value)
                              #and record resulting value
                              cgv[ind].append(phenotype_value)
               #print(cgv)
               # then calculate the population mean and s.d. over each condition
               cond_mean = np.mean(cgv, axis=0)
               cond_std = np.std(cgv, axis=0)


               #print(cond_mean)
               x = []
               y = []
               pop_mean = []
               env_cond = []

               for cond in range(0,env_range):
                      #print("environment: ", cond, "mean: ", cond_mean[cond], "s.d.: ", cond_std[cond])
                      dist_to_opt = abs(cond - sel_opt)
                      pop_mean.append(cond_mean[cond])
                      env_cond.append(cond)
                      x.append(dist_to_opt)
                      y.append(cond_std[cond])

               #plot population response
               #plt.scatter(env_cond, pop_mean, label='phenotypic population response')
               # Plot the estimated line
               #y_est = slope * x + intercept
               #plt.plot(x, y_est, label=f'Estimated line (slope={slope:.2f})')
               # Add labels and a legend
               #plt.xlabel('enviromental condition')
               #plt.ylabel('population mean')
               #plt.hlines(sel_opt, xmin = 0, xmax = env_range)
               #plt.vlines(sel_opt, ymin = 0, ymax = max(pop_mean))
               #plt.legend()
               # Show the plot
               #plt.show()


               slope, intercept, r_value, p_value, std_err = linregress(x,y)
               print("replicate:", reps)
               print("slope: ", slope, "r: ", r_value, "p :", p_value)
               print("selective optimum: ", sel_opt)
               print("mean phenotypic population distance to selective optimum", pop_mean[sel_opt] - sel_opt)
               print("percent deviation", pop_mean[sel_opt]/sel_opt)
               outf.write(str(replicates) + "\t" + str(slope) + "\t" + str(r_value) + "\t" + str(p_value) + "\t" + str(sel_opt) + "\t" + str(pop_mean[sel_opt] - sel_opt) + "\t" + str(pop_mean[sel_opt]/sel_opt) + "\n")

               # Plot the data points
               #plt.scatter(x, y, label='stdandard deviation')
               # Plot the estimated line
               #y_est = slope * x + intercept
               #plt.plot(x, y_est, label=f'Estimated line (slope={slope:.2f})')
               # Add labels and a legend
               #plt.xlabel('deviation from selective optimum')
               #plt.ylabel('standard deviation')
               #plt.legend()
               # Show the plot
               #plt.show()
```
